# Genome-wide fitness analysis identifies genes required for *in vitro* growth and macrophage infection by African and Global Epidemic pathovariants of *Salmonella* Enteritidis

**DOI:** 10.1101/2022.04.06.487138

**Authors:** Wai Yee Fong, Rocío Canals, Alexander V. Predeus, Blanca Perez-Sepulveda, Nicolas Wenner, Lizeth Lacharme-Lora, Nicholas Feasey, Paul Wigley, Jay C. D. Hinton

## Abstract

*Salmonella* Enteritidis is the second most common serovar associated with invasive non-typhoidal *Salmonella* (iNTS) disease in sub-Saharan Africa. Previously, genomic and phylogenetic characterisation of *S*. Enteritidis isolates from human bloodstream led to the discovery of the Central/Eastern African (CEAC) and West African clades, which were distinct from the gastroenteritis-associated Global Epidemic clade (GEC). The African *S*. Enteritidis clades have unique genetic signatures that include genomic degradation, novel prophage repertoires and multi-drug resistance, but the molecular basis for the enhanced propensity of African *S*. Enteritidis to cause bloodstream infection is poorly understood. We used transposon insertion sequencing (TIS) to identify the genetic determinants of the GEC representative strain P125109 and the CEAC representative strain D7795 for growth in three *in vitro* conditions (LB or minimal NonSPI2 and InSPI2 growth media), and for survival and replication in RAW 264.7 murine macrophages. We identified 207 *in vitro*-required genes that were common to both *S*. Enteritidis strains and also required by *S*. Typhimurium, *S*. Typhi and *Escherichia coli*, and 63 genes that were only required by individual *S*. Enteritidis strains. Similar types of genes were required by both P125109 and D7795 for optimal growth in particular media. Screening the transposon libraries during macrophage infection identified 177 P125109 and 201 D7795 genes that contribute to bacterial survival and replication in mammalian cells. The majority of these genes have proven roles in *Salmonella* virulence. Our analysis also revealed candidate strain-specific macrophage fitness genes, some of which represent potential novel *Salmonella* virulence factors.

**IMPACT STATEMENT:** Invasive non-typhoidal *Salmonella* (iNTS) disease is a systemic infection that has a high case fatality rate of 15% and is responsible for an estimated 66,500 deaths/year in sub-Saharan Africa. The main causative agents are pathovariants of *Salmonella* Typhimurium, known as *S*. Typhimurium ST313, and *Salmonella* Enteritidis (*S*. Enteritidis), known as Central/Eastern African (CEAC) and West African *S*. Enteritidis. Whilst the African *S*. Typhimurium pathovariant has been an active focus of research over the past decade, studies on African *S*. Enteritidis have been lacking. We used transposon insertion sequencing (TIS) to identify the genetic requirements of both African and Global Epidemic *S*. Enteritidis to grow *in vitro* and to infect murine macrophages. To our knowledge, this is the first genome-wide functional analysis of African *S*. Enteritidis under conditions relevant to infection of a mammalian host. We show that the gene sets required for growth under laboratory conditions and macrophage infection by African and Global Epidemic *S*. Enteritidis were broadly similar, and that the majority of the genes that contribute to survival and replication in macrophage already have proven roles in *Salmonella* virulence. Our analysis did identify candidate strain-specific macrophage fitness genes, some of which could be novel *Salmonella* virulence factors.

## INTRODUCTION

The majority of human pathogenic *Salmonella* belong to *S. enterica* subspecies 1, including the human-restricted serovars *S*. Typhi and *S*. Paratyphi (causative agents of typhoid and paratyphoid fever) and host-generalists *S*. Typhimurium and *S*. Enteritidis commonly associated with gastroenteritis infections. In recent years, non-typhoidal *Salmonella* (NTS) have emerged as the most common cause of community-onset bloodstream infections in sub-Saharan Africa [1, 2]. This invasive non-typhoidal *Salmonella* disease (iNTS) manifests as a febrile systemic illness resembling enteric fever that often lacks gastrointestinal symptoms, and disproportionately affects young children (under the age of five) with co-morbidities such as malnutrition, malaria or HIV infection or HIV-infected adults [3–6]. In 2017, iNTS disease was responsible for 77,500 deaths globally, of which 66,500 deaths occurred in sub-Saharan Africa [6]. The high case fatality-rate of iNTS (15%) [7] makes the disease a major health problem.

Most cases of human iNTS infections across Africa are caused by multi-drug resistant *S*. Typhimurium or *S*. Enteritidis variants that have characteristic genetic signatures that differ from gastroenteritis-associated *Salmonella* [1, 2, 8, 9]. African invasive *S*. Typhimurium isolates typically belong to the novel MLST sequence type 313 (ST313), which is distinct from the ST19 type that includes most gastroenteritis-associated *S*. Typhimurium isolates [8, 10]. A similar theme of phylogeographic differences has been observed in the *S*. Enteritidis serovar: two separate clades of African invasive *S*. Enteritidis have been identified, designated as the Central/Eastern African (CEAC) and West African clades, as opposed to the gastroenteritis-associated Global Epidemic clade (CEC) of *S*. Enteritidis [9]. Overall, the specific genomic signatures of African invasive *Salmonella* included multi-drug resistance determinants, distinct prophage repertoires and distinct patterns of genome degradation [8, 9, 11].

*S*. Enteritidis GEC strain P125109 and CEAC strain D7795 have been defined as the key representative strains of the respective clades. P125109 was isolated from an outbreak of human food poisoning in the United Kingdom in 1988 [12–14], while D7795 was isolated from blood culture from a Malawian child in 2000 [9]. The genome sequences of P125109 and D7795 were published in 2008 [12] and 2016 [9], respectively, and were recently reannotated and improved with long-read sequencing [15].

To date, genome-wide functional analyses of *S*. Enteritidis focused on gastroenteritis-associated *S*. Enteritidis in several *in vitro* conditions as well as during interaction with human epithelial cells, avian macrophages, and an animal infection model [16–19]. Other approaches investigated the role of specific genomic regions such as *Salmonella* Pathogenicity Islands (SPIs) and Regions of Difference (RODs) [20–23]. However, there have been no previous functional genomic analyses of African *S*. Enteritidis. The Feasey *et al*. [9] study remains the sole study to have examined and compared the metabolic capabilities and virulence of African and Global Epidemic *S*. Enteritidis using a variety of carbon sources and an avian infection model. Because the primary intracellular niche of *Salmonella* during systemic infection, however, is macrophages [24], it was important to generate a comprehensive description of *S*. Enteritidis genes required for macrophage infection. To gain insights into how *S*. Enteritidis CEAC D7795 causes disease, it was important to determine whether this pathovariant carries novel virulence genes that have not been described previously in GEC P125109 or other *Salmonella* serovars.

Here, we used transposon insertion sequencing (TIS) to characterise the gene functions of the *S*. Enteritidis GEC P125109 and CEAC D7795 genomes in three different *in vitro* growth conditions and during macrophage infection (Fig 1). TIS involves the random transposon insertional mutagenesis and high-throughput sequencing of the *S*. Enteritidis genomes following selection under different experimental conditions. The relative changes in abundance of each transposon mutant before and after the treatment reflects the contribution of each gene to survival and adaptation to a particular environment. TIS has been successfully used in various bacteria species to define genetic requirements for bacterial viability and fitness during *in vitro* growth and following infection of mammalian hosts and host cells [25–33]. Our findings delineate the fitness landscapes of GEC P125109 and CEAC D7795 during conditions relevant to mammalian host infection. We identified key similarities and differences in gene requirements between the representatives of the two *S*. Enteritidis clades, coupled with extensive correlation to other *Salmonella* serovars.

**Fig 1.**
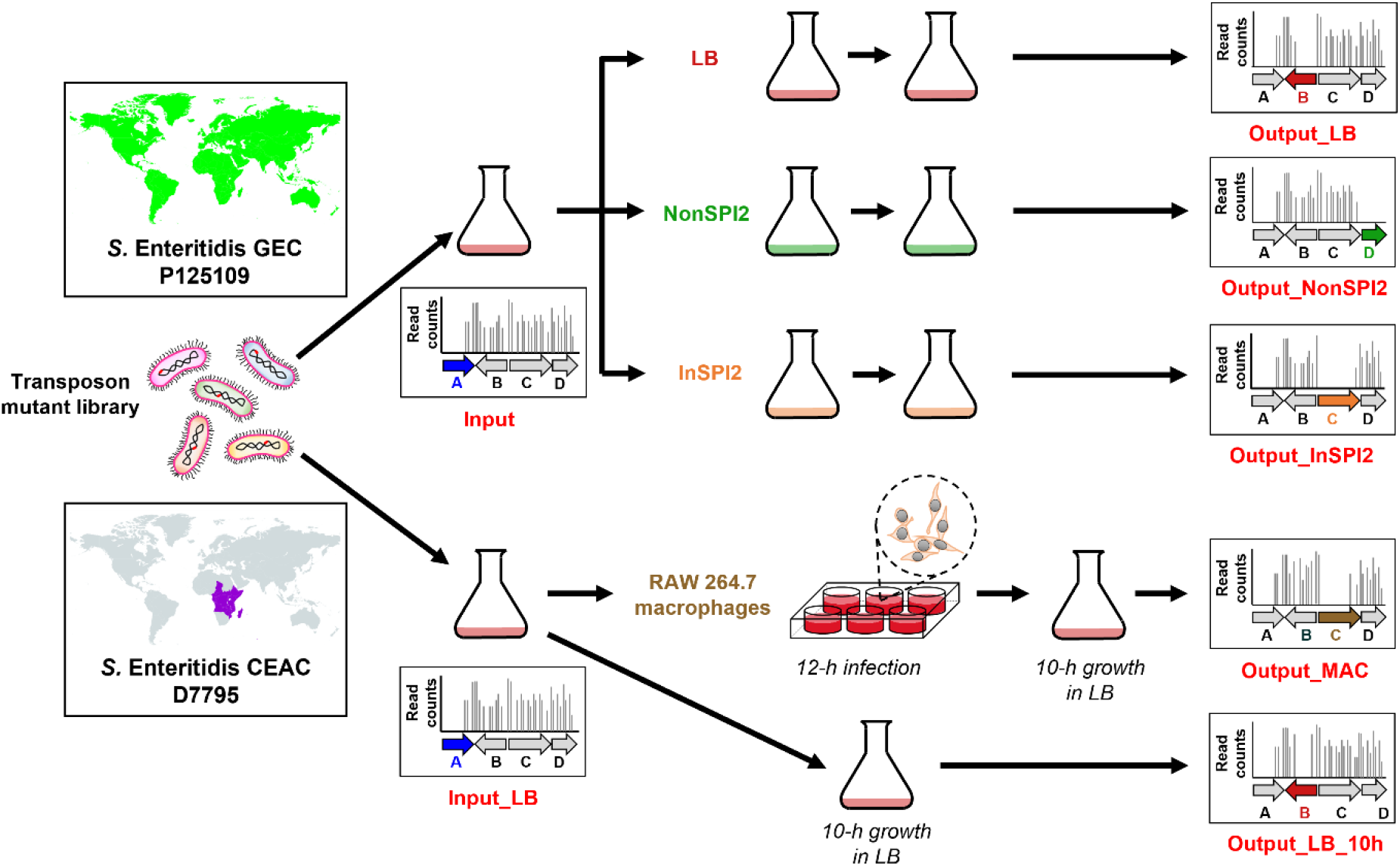
Transposon-insertion sequencing (TIS) of both *S*. Enteritidis Global Epidemic clade (GEC) strain P125109 and Central/Eastern African clade (CEAC) strain D7795. Schematic representation of the *S*. Enteritidis transposon libraries and growth conditions used in this study. Experimental details are provided in Materials and Methods. The genes (*A*, *B*, *C* or *D*) highlighted with colour in the five right-hand panels illustrate how required or fitness genes for a particular environmental condition were identified.

## MATERIALS AND METHODS

### Bacterial strains and growth conditions

Bacterial strains used in this study are shown in Table S1. Permission to work with the *S*. Enteritidis CEAC strain D7795 was approved by the University of Malawi College of Medicine Research Ethics Committee (COMREC ethics no. P.06/20/3071).

The Lennox formulation of Lysogeny Broth, (LB) was 10 g/L tryptone (Difco), 5 g/L yeast extract (Difco), and 5 g/L NaCl (Sigma). LB agar was prepared by addition of 15 g/L Bacto Agar to LB media prior to autoclaving. InSPI2 (pH 5.8, 0.4 mM inorganic phosphate [P_i_]) and NonSPI2 (pH 7.4, 25mM Pi) media are PCN-based synthetic minimal media [34, 35]. Terrific broth (TB) (Sigma), a nutrient-rich medium for higher growth of bacteria [36], was prepared according to the manufacturer’s instructions. SOC media contained 20g/L tryptone, 5 g/L yeast extract, 0.5 g/L NaCl, 2.5 mM KCl, 10 mM MgCl_2_ and 20 mM glucose [37].

Bacterial cultures were inoculated from a single colony routinely and grown in 5 mL LB broth in a sterile 30 mL screw-capped glass vial, for 16 h at 37°C with shaking at 220 rpm (this procedure is referred to as ‘overnight culture’ unless otherwise described). For experiments, bacterial cultures were grown in 25 mL media in 250 mL sterilised Erlenmeyer flasks, unless otherwise indicated. When required, the antibiotic kanamycin (Km) was added to a final concentration of 50 μg/mL, tetracycline (Tet) to 25 μg/mL, and gentamicin (Gm) to 20 μg/mL.

### Preparation of electro-competent cells and electroporation

The electroporation of *S*. Enteritidis was carried out using an adaptation of the method of Dower *et al*. [38].

For the preparation of electro-competent cells, overnight cultures were diluted 1:100 into 25 mL salt-free LB (unless otherwise indicated) and grown at 37°C (or 30°C for strains carrying temperature-sensitive plasmids) at 220 rpm to OD_600_ 0.45 ± 0.05. The appropriate antibiotic was added for plasmid-carrying strains. For strains harbouring the λ Red recombination plasmid pKD46, L-arabinose was added to a final concentration of 0.2%. Cells carrying the pSIM5-*tet* plasmid were grown at 30°C to OD_600_ of 0.40 ± 0.05 and then incubated at 42°C for 15 min with shaking, to stimulate the expression of the λ Red recombinase.

At the desired OD_600_, bacterial cells were transferred to 50 mL centrifuge tubes, chilled on ice for 10 min, then pelleted by centrifugation at 4°C and 4,000 rpm for 10 min. Cells were washed three times in 25 mL ice-cold sterile H_2_O, and finally resuspended in 250 μL of ice-cold sterile 10% (v/v) glycerol. Cells were aliquoted into 50 μL volumes for electroporation reactions, or storage at −80°C.

For electroporation, competent cells were mixed with 500 ng of DNA in electroporation cuvettes (2 mm gap) and the reactions were electroporated (2.5 kV) using a MicroPulser electroporator (Bio-Rad). Electroporated cells were recovered in 1 mL LB (unless otherwise stated) at 37°C for 1 h (or 30°C for 2 h if the cells contained temperature-sensitive plasmids) with shaking at 220 rpm, before plating on LB agar plates containing the appropriate antibiotic to select for transformants.

### Construction of *S*. Enteritidis transposon mutant library

Libraries of transposon insertion mutants were constructed in *S*. Enteritidis strains P125109 and D7795 using the EZ-Tn5™ <KAN-2> Insertion Kit (Lucigen) as previously described [31]. Briefly, transposome mixtures were prepared by mixing glycerol, TypeOne™ Restriction Inhibitor (Lucigen), EZ-Tn5<KAN-2> transposon (at 0.1 pmol/μL) and EZ-Tn5 Transposase, and electroporated into P125109 or D7795 competent cells. Electroporated cells were grown in SOC media for 1 h, before plating on multiple LB agar plates containing 50 μg/mL Km followed by overnight incubation at 37°C to select transformants. Following colony counting, the transposon mutants were collected from the plates by adding LB liquid media, and pooling together for growth in LB + Km^50^ at 37°C overnight to generate the transposon mutant library.

### Construction of mutants in *S*. Enteritidis by λ Red recombineering

Mutants were constructed using the λ Red recombination method [39]. The oligonucleotides phoQ_KO_F and phoPQ_KO_R were used to amplify the Km resistance cassette from the pKD4 plasmid. The polymerase chain reaction (PCR) product was electroporated into P125109 cells containing the pSIM5-*tet* plasmid [40] and D7795 cells containing the pKD46-*aacC1* plasmid, respectively, to replace the *phoPQ* genes. Transformants were selected on LB agar plates containing 25 μg/mL Tet or 20 μg/mL Gm at 37°C. Colony PCR was used to confirm the presence of the deletion mutation in transformant colonies.

To generate non-polar, in-frame deletions of *phoPQ*, the corresponding Km-resistant derivatives were transformed with the temperature-sensitive pCP20-*tet* [41] (for P125109) or pCP20-Gm [42] (for D7795) plasmid that synthesises the FLP recombinase. Transformants were selected by overnight growth at 30°C on LB agar plates containing Tet^25^ or Gm^20^, followed by passaging on LB agar plates at 37°C to cure the pCP20 plasmid. Loss of the antibiotic resistance cassette and pCP20 was confirmed by checking for loss of resistance to Km and Tet (or Gm). Presence of the gene deletion was also confirmed by colony PCR.

Electro-competent cells of wild-type P125109 and D7795, and P125109 derivatives were prepared following growth in LB media as described earlier. Because of the relatively low electroporation efficiency of the D7795 derivative strains following growth in LB, electro-competent cells of D7795 containing pKD46 and Km-resistant derivatives were prepared following growth in TB media.

The construction of *Salmonella* mutants by FRT-mediated gene deletion generally involves a subsequent P22 bacteriophage transduction into a clean wild-type background. However, observations from P22 plaque assays performed with P125109 and D7795 (data not shown) indicated that P22 did not infect these strains readily. To ensure that no unintended nucleotide changes had been generated by the λ Red mutagenesis process, one *phoPQ* deletion mutant per strain was whole-genome sequenced (MicrobesNG, Birmingham, UK). Bioinformatic analysis of the sequencing data involved comparison with the genome of either P125109 and D7795 [15], which confirmed that no unintended mutations had been introduced during the recombineering process (data not shown).

### Passaging the *S*. Enteritidis transposon libraries in LB, NonSPI2 and InSPI2 media

A 1.5 mL aliquot of the P125109 or D7795 transposon library was grown in 25 mL LB + Km^50^ in a shaking water bath at 37°C, 220 rpm for 16 h. Cells harvested from 4 x 200 μL aliquots of the bacterial overnight culture were stored at −80°C prior to genomic DNA extraction, to give the Input sample. Another 1 mL of the bacterial culture was washed twice with phosphate buffered saline (PBS) and resuspended in LB, NonSPI2 or InSPI2 media. A 1:100 dilution was inoculated into 25 mL of LB, NonSPI2 or InSPI2 media (without antibiotic), respectively, in 250 mL Erlenmeyer flasks. Cultures were incubated in a shaking water bath at 37°C, 220 rpm until early stationary phase (ESP) (passage 1). ESP was defined as the moment when the growth of the bacterial culture in the respective growth media first reached a plateau, as measured by OD600 readings. The ESP timepoints for P125109 were 7 h (LB), 10 h (NonSPI2), 10 h (InSPI2) and for D7795 were 6 h (LB), 24 h (NonSPI2), 24 h (InSPI2).

A total of two passages were performed in each growth medium. For LB passages, 250 μL culture was transferred in each individual passage, following two washes with PBS. For NonSPI2 and InSPI2 passages, to account for the reduced growth that occurred in the minimal medium, a culture volume with OD_600_ equivalent to the LB subculture inoculum was transferred into the subsequent passage to make bacterial numbers as equivalent as possible. For example, if LB passage 1 has an OD_600_ of 2.0 and NonSPI2 passage 1 has an OD_600_ of 1.0, then 500 μL of NonSPI2 passage 1 was used to inoculate passage 2. Cells from 4 x 200 μL aliquots of the second passage of LB (Output_LB), NonSPI2 (Output_NonSPI2) and InSPI2 (Output_InSPI2) were harvested and stored at −80°C until genomic DNA extraction.

### Infection of RAW 264.7 macrophages with *Salmonella*

For intra-macrophage replication assays with the wild-type and del-*phoPQ* derivatives of *S*. Enteritidis and *S*. Typhimurium strains, 10^6^ RAW 264.7 macrophage cells (ATCC® TIB-71™) were seeded in each well of 6-well plates (Sarstedt) 24 h prior to infection. Bacterial overnight cultures were inoculated from a single bacterial colony into 25 mL LB, incubated shaking at 220 rpm and 37°C for 18 h. Inoculum size was standardised by adjusting the OD_600_ of overnight cultures to OD_600_ 2.0, followed by resuspension in Dulbecco’s Modified Eagle Medium (DMEM; Thermo Fisher Scientific) supplemented with MEM non-essential amino acids (NEAA) (Thermo Fisher Scientific; 10% final concentration) and L-glutamine (Thermo Fisher Scientific; 2 mM final concentration). Prior to all macrophage infection experiments, bacteria were opsonised with 10% BALB/c mouse serum (Charles River) in 10 volumes of DMEM for 30 min on ice.

The macrophages were infected with *Salmonella* at a Multiplicity of Infection (MOI) of 5–10, and infections were synchronised by 5 min centrifugation at 1,000 rpm at room temperature. This was defined as time 0. After 30 min incubation at 37°C and 5% CO_2_, cells were washed three times with Dulbecco’s phosphate-buffered saline (DPBS) and incubated with DMEM + 10% foetal bovine serum (FBS; Thermo Fisher Scientific) containing 100 μg/mL Gm for 1 h to kill extracellular bacteria. For time points beyond 1.5 h post-infection, the cell culture media was replaced with fresh DMEM + 10% FBS containing 10 μg/mL Gm. Intracellular bacterial numbers were determined by lysis of infected macrophages at 1.5 h and 15.5 h post-infection with 1% Triton X-100 (in DPBS). Serial dilutions of the cell lysates were plated onto LB agar plates (containing antibiotics where necessary) and incubated overnight at 37°C for bacterial enumeration. Replication fold-change was calculated using the intracellular numbers at 15.5 h vs. 1.5 h.

For macrophage infection with the *S*. Enteritidis P125109 and D7795 transposon libraries, 10^6^ RAW 264.7 macrophages were seeded in each well of 6-well plates 24 h before infection. A 1.5 mL aliquot of P125109 or D7795 transposon library was grown in 25 mL LB + Km^50^ in a shaking water bath at 37°C, 220 rpm for 16 h, and genomic DNA was isolated from two different biological replicates as input samples (Input_LB_1 and Input_LB_2). OD_600_ of the overnight culture was measured and a bacterial inoculum equivalent to OD_600_ = 5.0 was prepared by pelleting and resuspending bacterial cells in appropriate volumes of DMEM supplemented with MEM NEAA and L-glutamine; this equilibration step ensured that sufficient bacterial cells (~1.4 x 10^7^ cells for P125109 and ~1.5 x 10^7^ cells for D7795) were used in infection to represent the complexity of the transposon library. Macrophages were infected at an MOI of 5–10 with mouse serum-opsonised bacteria, as described earlier. A total of 18 wells were used in each infection per strain, with 6 wells set aside for the generation of bacterial counts at 1.5 h and 12 h post-infection, and calculation of the fold-change replication of the intracellular bacteria (12 h vs. 1.5 h).

At 12 h post-infection, macrophages were lysed with 1% Triton X-100. Macrophage lysates containing intracellular bacteria from 12 wells were pooled into one 15 mL centrifuge tube and centrifuged at 4,000 rpm for 5 min. The cell pellet was resuspended in 1 mL LB and transferred to a flask containing 24 mL LB supplemented with Km^50^ for 10 h growth at 37°C, 220 rpm (Output_MAC). To determine the effect of 10 h growth in LB on the transposon library, a fraction of the input library culture was sub-cultured in LB containing Km^50^ for 10 h (Output_LB_10h). Cells were harvested from the output transposon library cultures and stored at −80°C until genomic DNA extraction.

### DNA manipulation and sequencing

Genomic DNA was purified from all input and output library cultures using the Quick-DNA™ Miniprep Plus Kit (Zymo Research), following the manufacturer’s instructions. To ensure that sufficient genomic DNA was available for the preparation of Illumina DNA libraries, each DNA sample comprised DNA extracted from 4 x 200 μL aliquots of the respective bacterial cell samples. DNA concentrations (in ng/μL) were determined using the Qubit dsDNA High Sensitivity Assay and the NanoDrop 2000 spectrophotometer (Thermo Fisher Scientific).

For Illumina DNA library preparation, 2 μg of genomic DNA from each mutant pool was first fragmented to an average size of 300–350 bp using the BioRuptor@Pico sonication system (15 s ON 90 s OFF, 9 cycles). Illumina DNA library preparation was performed using NEBNext® DNA Library Prep Master Mix Set for Illumina® (New England Biolabs), following the manufacturer’s instructions. Reaction products from each step of library preparation were purified using AMPure XP beads (Beckman Coulter).

To amplify the transposon-flanking regions, transposon-specific forward oligonucleotides (Table S1) were designed such that the first 10 bases of each Read 1 (R1) would be the transposon sequence. A unique 6-base barcode was incorporated into the forward oligonucleotide to allow the pooling of samples for multiplex sequencing in a single lane. 22 cycles of PCR [26, 31] were performed with NEBNext Q5 Hot Start HiFi polymerase using the transposon-specific oligonucleotides and the Illumina reverse primer PE PCR Primer 2.0 for each fragmented DNA sample, following the recommended denaturation, annealing and extension temperatures and durations for NEBNext Q5 Hot Start HiFi polymerase. The resulting DNA was quantified using Qubit dsDNA High Sensitivity Assay (Thermo Fisher Scientific) and visualised on an Agilent High Sensitivity DNA chip (Agilent Technologies), following the manufacturer’s instructions. Finally, the amplified library was purified with AMPure XP beads and eluted in 30 μL of molecular grade H_2_O.

The list of Illumina DNA libraries generated in this study is given in Table S2. For sequencing, the Illumina DNA libraries from P125109 and D7795 were pooled in a ratio corresponding to the difference in estimated library complexity (which was initially defined by the number of transformant colonies) between the two strains. The DNA libraries from RAW 264.7 macrophage infection experiments were pooled in the ratio of 3:1 (P125109:D7795). The DNA libraries from *in vitro* passages in LB, NonSPI2, InSPI2 were pooled in the ratio of 2:1 (P125109:D7795); the ratio was revised following sequencing of the DNA libraries from macrophage infection experiments, where the actual transposon library densities were revealed to be ~200,000 unique insertions for P125109 and ~100,000 insertions for D7795.

QC assessment of the pooled DNA library and sequencing were performed by the Centre for Genomic Research (CGR), University of Liverpool. Each library pool was size-selected to 250–500 bp, then paired-end sequenced in one lane on an Illumina HiSeq4000 at 2 x 150 bp (for DNA libraries generated from macrophage infection experiments) or two lanes on an Illumina NovaSeq6000 (SP mode) at 2 x 150 bp (for DNA libraries generated from *in vitro* passage experiments) respectively. 15% of the bacteriophage ϕX174 genome, provided by Illumina as a control, was added to each lane to overcome the low complexity of the bases that followed the barcode in R1 [43].

### *S*. Enteritidis genome sequences and annotations

The annotated complete long-read-based genome assemblies of *S*. Enteritidis P125109 and D7795 are available in the National Center for Biotechnology Information (NCBI) Assembly database (accession numbers SAMN16552335GCA_015240635.1 [P125109] and [SAMN16552336GCA_015240855.1](https://www.ncbi.nlm.nih.gov/assembly/GCA_015240635.1) [D7795]) [15]. Orthologues between the *S*. Enteritidis and other *Salmonella* strains presented in this study were identified using the pipeline described at https://github.com/apredeus/multi-bacpipe. Briefly, Roary [44] was used to find protein-coding orthologues, followed by nucleotide BLAST (blastn) to find conserved non-coding sRNAs, genes encoding small proteins, and pseudogenes. Final validation of each annotation was achieved manually by comparing the annotations to the locus tags in the published P125109 annotation. Cluster of Orthologous Genes (COG) categories were assigned with eggNOG-mapper v2 [45, 46] using the default parameters.

### Sequence analyses of the *S*. Enteritidis transposon library

Bioinformatic processing and analysis of *S*. Enteritidis transposon insertion data followed the general strategy described in [26, 31, 43], with modifications. The code and full description of the pipeline are available at https://github.com/apredeus/TRADIS.

Raw sequencing data was demultiplexed using cutadapt v2.6 [47]. First, a barcode sequence fasta file that included one sequence per sample was compiled, after which cutadapt was run with options “cutadapt -O 34 -g file:barcodes.fa --discard-untrimmed”. This generated a set of two paired-end fastq files for each sample. The reads were then aligned to the reference genome using bwa v0.7.17-r1188 (using bwa mem algorithm). Aligned BAM files were sorted and indexed using samtools v1.9 [48].

For further processing, two GFF annotation files were generated for each bacterial strain used in the experiments. One file was used for deduplicated read counting and was obtained from a general annotation file by changing the type of each feature that had a locus tag (ID) into “gene”. The second GFF file was generated the same way, with additional change to the annotated feature size: the last 10% of each annotated gene was removed. This annotation file was used in essentiality analysis, since previous reports [43, 49] showed that insertions or deletions in the last 10% of the gene are much less likely to cause a complete loss of function.

Following these steps, a series of additionally processed alignment (BAM) files was created. First, picard MarkDuplicates v2.21.2 was used to remove PCR and optical duplicates from the aligned reads. The resulting files were filtered using *cigar_filter.pl* to select only the reads that align exactly at the start of R1 without softclipping. The resulting filtered BAM files were converted into 1 nucleotide single-end BAM files using *make_1nt.pl* script to avoid counting reads that spanned into the nearby genes. These alignment files were used for quantification of deduplicated reads and used in DESeq2 analysis. In parallel, the filtered BAM files were also converted into 1 nt unique insertion BAM files using *make_1nt_uniq.pl* script; these files were used for essentiality analysis.

The resulting 1 nt BAM files were quantified using featureCounts v1.6.4 [50] with “-M -O --fraction -t gene -g ID -s 0” options for DESeq2 analysis, and “-M -O -t gene -g ID -s 0“ options for essentiality analysis. This was done to account for multimapping reads: if only uniquely mapping reads were considered, transposable and other repetitive elements looked falsely essential. Indeed, if multimapping reads were discarded, genes that have multiple copies (such as transposons) appear to have a zero transposon insertion rate, which in turn leads us to the false conclusion of their essentiality.

### Essentiality analysis

Essentiality analysis was done using the unique insertion counts. For consistency with our DESeq2-based fitness analysis, we have re-implemented the essentiality analysis functions of Bio-Tradis [43], specifically R functions *make_ess_table* and *calculate_essentiality*. Briefly, the unique insertion counts were converted into an insertion index (that is, insertion sites divided by gene length) [26], which followed the expected bimodal distribution. The distribution histogram was used to fit two functions: exponential function for very low insertion indices corresponding to the required genes, and gamma function for high insertion indices of the dispensable genes. Using the obtained fits, the genes were classified as follows: if log-likelihood of the exponent to gamma distribution was 2 or higher, the gene was deemed “required”; if the ratio was below −2, it was deemed “not required”; genes with intermediate insertion indices were reported as “ambiguous”. Due to the relatively low insertion densities in our libraries, essentiality calls were not assigned for features shorter than 200 nt.

Data from the TIS-based study on *S*. Typhimurium ST313 D23580 [31] was used as a comparator to identify common *Salmonella* genes that were required for growth under laboratory conditions or during macrophage infection, and differences that might reflect unique requirements for each serovar or the pathogenic niche inhabited by these bacteria. Due to the substantial differences in the average number of unique insertions present in the transposon library for *S*. Enteritidis GEC P125109, CEAC D7795 and *S*. Typhimurium D23580, we used both deduplicated read counts and essentiality calls to identify robust differences.

Deduplicated read counts for LB input, LB output, and macrophage output for the three strains were log_2_-transformed and quantile-normalised. These values were then used to calculate a log_2_ fold-change between the libraries that was evaluated and found to follow an approximately normal distribution (data not shown). Thus, we have selected the genes that were significantly different in all three conditions in the particular strains according to t-test (*P*-value ≤ 0.05), and also had differing essentiality calls. This approach allowed us to identify 63 genes that satisfied both conditions. The resulting genes were visualised using Phantasus gene expression analysis tool (http://genome.ifmo.ru/phantasus-dev/). K-means clustering of rows allowed us to identify five distinct groups of genes, according to the strain they were required in (Fig S1).

### DESeq2-based fitness analysis

Analysis of differential fitness was performed in R v4.0.2, using DEseq2 v1.28.1 [51] and a simple study design (~ Condition) with default settings. Deduplicated raw read counts were used as expression values. Each condition was represented by at least two biological replicates. Results are shown in log_2_ fold-change. A cut-off of 2-fold-change and *P*-value < 0.05 was applied.

### Statistical analyses of intra-macrophage replication experiments

Statistical analyses were performed with GraphPad Prism 7 (version 7.04). Ordinary one-way ANOVA and the Bonferroni’s multiple comparison test were used to determine differences in intra-macrophage replication levels between different *Salmonella* strains. A *P*-value of less than 0.05 was considered to be statistically significant.

## RESULTS AND DISCUSSION

### Characterisation of the *S*. Enteritidis P125109 and D7795 transposon libraries

Transposon insertion libraries were constructed in the *S*. Enteritidis GEC representative strain P125109 and CEAC representative strain D7795. Each pool of transposon mutants was grown in LB (Input) and passaged two times successively at 37°C in three different growth media: a nutrient rich medium, LB (Output_LB), an acidic phosphate-limiting PCN-based minimal media that induces SPI-2 expression, designated InSPI2 (Output_InSPI2), and a neutral pH PCN-based minimal media that does not induce SPI-2 expression, designated NonSPI2 (Output_NonSPI2) [34] (Fig 1).

Genomic DNA from the input and output samples was purified and prepared for high-throughput Illumina sequencing of the DNA adjacent to the transposon. Sequencing was performed on two lanes of an Illumina NovaSeq6000 system, generating a total of 1,048,450,846 paired-end reads (523,450,580 read pairs from the first lane and 525,000,266 read pairs from the second lane). The sequencing data were processed as described in Materials and Methods. Table S2 shows the number of sequence reads obtained, the sequenced reads that contained the transposon tag sequence, and the sequence reads that were mapped to the respective *S*. Enteritidis genomes.

Sequence analyses of the input libraries identified 246,743 unique transposon insertion sites in P125109 and 195,646 unique insertions in D7795, an average of one insertion per 19 nucleotides in the P125109 genome and one insertion per 24 nucleotides in the D7795 genome. The complete data sets are available for visualisation in two online JBrowse genome browsers for the respective strains: https://tinyurl.com/GECP125109 and https://tinyurl.com/CEACD7795. The browsers show the transposon insertion profiles for the chromosome and plasmid of *S*. Enteritidis (pSENV in P125109; pSEN-BT and pRGI00316 in D7795). To yield maximum biological insight from the transposon mutagenesis experiment, we employed our recent re-annotation of the coding genes and non-coding sRNA genes of P125109 and D7795 [15], which was derived from a comparative genomic approach that identified all the annotated genes of *S*. Typhimurium ST19 [52] and *S*. Typhimurium ST313 [53] that were carried by the two *S*. Enteritidis strains (Materials and Methods).

An insertion index was calculated for each gene by dividing the number of unique insertions for any given gene by gene length; the data were used for essentiality analyses (Materials and Methods), classifying genes as “required, “not required” and “ambiguous”. “Required” genes in this study included genes essential for bacterial viability (i.e. genes that when disrupted lead to irreversible growth arrest or cell death), and genes that contribute strongly to fitness in a particular environmental condition [31, 54]. The role of genes shorter than 200 nt in length could not be defined robustly and was designated as “short”. The number of reads, transposon insertion sites, insertion index, and the essentiality calls per gene for all conditions tested are summarised in Table S3.

### Identification of *S*. Enteritidis P125109 and D7795 genes required for *in vitro* growth

Essentiality analysis of the *S*. Enteritidis GEC strain P125109 input library identified 497 required genes, 3516 dispensable genes and 317 ambiguous genes, with 693 genes being classified as short (Fig 2). Of the 497 required genes (Table S3), 492 genes were located on the chromosome and 5 genes were carried by the pSENV virulence plasmid (*traJ*, *samA*, *samB*, *SEN_p0037* and *SEN_p0021*). To provide a functional context for the required genes, we used eggNOG-mapper v2 to assign Cluster of Orthologous Genes (COG) functional categories (Table S7). The majority of the required genes were involved in translation (J, 17%) and cell wall biogenesis (M, 5%), followed by an approximately equal distribution between the categories of transcription (K, 3%), replication (L, 3%), energy production (C, 3%) and various metabolic processes (E, F, H and I). Ninety-six genes (16%) were classified in the “Poorly Characterised” category, including 28 genes belonging to Regions of Difference (RODs) [12], prophage regions and SPIs. Eleven SPI-associated genes were required by P125109 for *in vitro* growth, including several *ssa* and *ttr* genes in SPI-2, *SEN0277* in SPI-6, and *SEN4250* in SPI-10.

**Fig 2.**
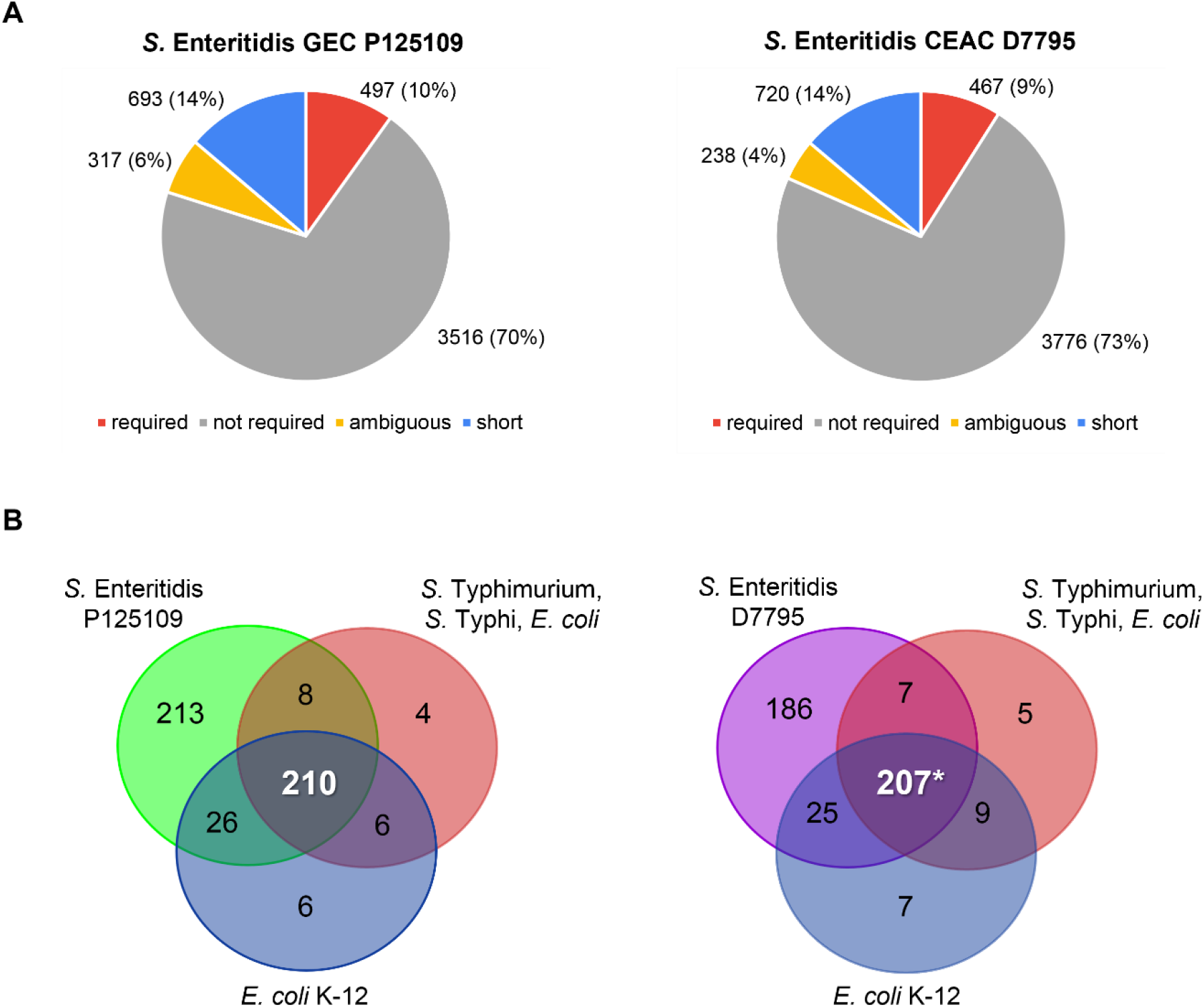
*S*. Enteritidis GEC P125109 and CEAC D7795 genes required for *in vitro* growth. (A) Distribution of *S*. Enteritidis GEC P125109 (left panel) and CEAC D7795 (right panel) genes into required, not required, ambiguous and short (gene length < 200 nt) categories. Full gene lists are presented in Table S3. (B) Comparison of required genes identified in GEC P125109 and CEAC D7795 with published gene essentiality studies for *S*. Typhimurium [25], *S*. Typhi [25] and *E. coli* [25, 49]. Only genes that shared an orthologue in P125109 or D7795 were used in the comparison. The asterisk (*) indicates that all the 207 D7795 genes were included in the 210 required P125109 genes (Table S4). Venn diagrams were generated using http://bioinformatics.psb.ugent.be/webtools/Venn/.

Essentiality analysis of the *S*. Enteritidis CEAC strain D7795 input library identified 467 required genes, 3776 dispensable genes, 238 ambiguous genes and 720 short genes (Fig 2). Of the 467 required genes (Table S3), 465 genes were encoded in the chromosome and 2 genes (*samA* and *SEN_p0046*) were located on the virulence plasmid pSEN-BT; no required genes were carried by the smaller plasmid pRGI00316. COG categories were assigned to the D7795 reference genome as described earlier. As seen for P125109, the majority of the D7795 required genes are involved in translation (J, 18%), cell wall biogenesis (M, 9%), and nucleotide metabolism (F, 7%). Nine required genes were associated with pathogenicity islands, namely four SPI-2 genes (*ssaT*, *ssaH*, *ttrAB*), two SPI-6 genes (*SEN2077* and *SEN2078*), two SPI-10 genes (*SEN4248* and *SEN4250*), and one SPI-14 gene (*SEN0801*).

### The 207 required genes shared between *S*. Enteritidis P125109 and D7795, and *S*. Typhimurium, *S*. Typhi and *E. coli*

To put our essentiality analysis into context, we drew upon other studies that used similar TIS approaches. The required genes of *S*. Enteritidis GEC P125109 and CEAC D7795 were compared with the genetic requirements of *Salmonella* serovars Typhimurium and Typhi, and *Escherichia coli*. We used two required gene sets: the first is a list of 228 essential genes shared between *S*. Typhimurium, *S*. Typhi and *E. coli* [25], and the second list of 248 essential genes identified in *E. coli* K-12 [49]. We found that a total of 207 genes were required in all *S*. Enteritidis, *S*. Typhimurium, *S*. Typhi and *E. coli* strains (Fig 2). These 207 required genes mainly encode the basic cellular machinery (e.g. DNA replication and protein translation) and pathways vital for the growth of the bacteria, including cell wall biogenesis and cell division (Table S4). Of the 6 and 9 genes identified as required in *S*. Typhimurium, *S*. Typhi and *E*. *coli* but not in P125109 or D7795, respectively, most were designated as “short” genes (gene length less than 200 nt) in either *S*. Enteritidis strain and therefore had not been assigned an essentiality call. Such genes included *infA*, encoding translation initiation factor IF-1; *csrA*, encoding a post-transcriptional regulator that regulates metabolism important for establishing infection in the intestine [55]; *rpmD*, *rpmC* and *rpmH*, encoding 50S ribosomal proteins. The *ispB* gene was the only one of these genes identified as not required in D7795. Overall, our observations were in agreement with other reports: required genes are conserved among bacterial strains within the same species or between different species [25, 27, 29, 31, 56].

### The 63 orthologous genes that are only required by *S*. Enteritidis or *S*. Typhimurium D23580

The Venn comparisons in Fig 2 suggest there are approximately 200 genes that are only required by *S*. Enteritidis but dispensable in *S*. Typhimurium, *S*. Typhi and *E. coli*. However, the differences that arise from such direct comparisons involving different TIS-based studies can be challenging to interpret, in part due to the differences in experimental protocols, transposon library profiles and bioinformatic processing pipelines used by different laboratories. Consequently, we made use of data from our recently published TIS-based analysis of genetic requirements of *S*. Typhimurium ST313 D23580 for survival and growth both *in vitro* and during macrophage infection [31] to identify differences in the genetic requirements with *S*. Enteritidis P125109 and D7795. To address the issue of the different insertion densities of the two *S*. Enteritidis transposon libraries (~200,000 each) and the *S*. Typhimurium ST313 D23580 library (at least ~500,000), a customised pipeline (Materials and Methods) was used to compare the three strains and to perform essentiality analysis. Both essentiality calls (using insertion indices) and changes in abundance of read counts in three conditions (LB input, LB output, and macrophage output) were used in the inter-strain essentiality analysis.

A total of 63 orthologous genes were identified as differentially required by *S*. Enteritidis P125109, D7795 and *S*. Typhimurium ST313 D23580, and broadly classified into five groups (Fig S1). All 63 genes are >200 nt, indicating that their essentiality calls are reliable. Identification of the chromosomal *cysS* gene as not required in *S*. Typhimurium ST313 D23580 provides support for our inter-strain essentiality analysis approach: a second orthologous *cysS* gene is encoded by the pBT1 plasmid of D23580, and we have previously shown experimentally that the chromosomal *cysS* gene is dispensable for growth [31].

Genes belonging to group 3 are of particular interest (Fig S1). Group 3 represents genes that are not required by the two African *Salmonella* strains D7795 and D23580, but are required by the GEC strain P125109. Pseudogenisation, and the consequent loss of gene function, is linked to a restriction in host range of *Salmonella*, as observed for *S*. Typhi [57] and for the switch from an enteric to an extra-intestinal lifestyle by African *S*. Typhimurium [11, 58]. For the future, it will be important to determine whether the functions of these 22 genes are truly dispensable for *S*. Enteritidis CEAC D7795 and *S*. Typhimurium ST313 D23580.

### Identification of fitness genes of *S*. Enteritidis P125109 and D7795 during growth in LB, NonSPI2 and InSPI2 *in vitro* conditions

To build upon our identification of required genes in P125109 and D7795, we studied *in vitro* fitness of the two *S*. Enteritidis strains by analysing the transposon mutants recovered after two passages in three different growth media under laboratory conditions: LB, NonSPI2 and InSPI2. The fitness genes required for growth in each media were identified by the insertion index and essentiality analysis (Table S3).

Following two passages of the P125109 transposon library in complex LB media, 538 genes were designated as required, which included genes that were required in the input as well as genes that contributed to fitness for *in vitro* growth in LB: 531 genes in the chromosome and 7 genes in the pSENV plasmid. A total of 565 genes were required for optimal growth after two passages in NonSPI2 minimal media: 561 genes in the chromosome and 4 genes in the pSENV plasmid. Analysis of the InSPI2 output library identified 629 genes, with 622 genes located in the chromosome and 7 genes in the pSENV plasmid. The overlap between the required gene lists for input, LB output, NonSPI2 output and InSPI2 output was determined to distinguish between the genes shared between all conditions and genes that were only required under specific conditions. We identified 19 genes for LB, 14 genes for NonSPI2, and 48 genes for InSPI2 that were required only in the specific media; these genes were categorised as LB-only, NonSPI2-only and InSPI2-only, respectively (Fig 3).

**Fig 3.**
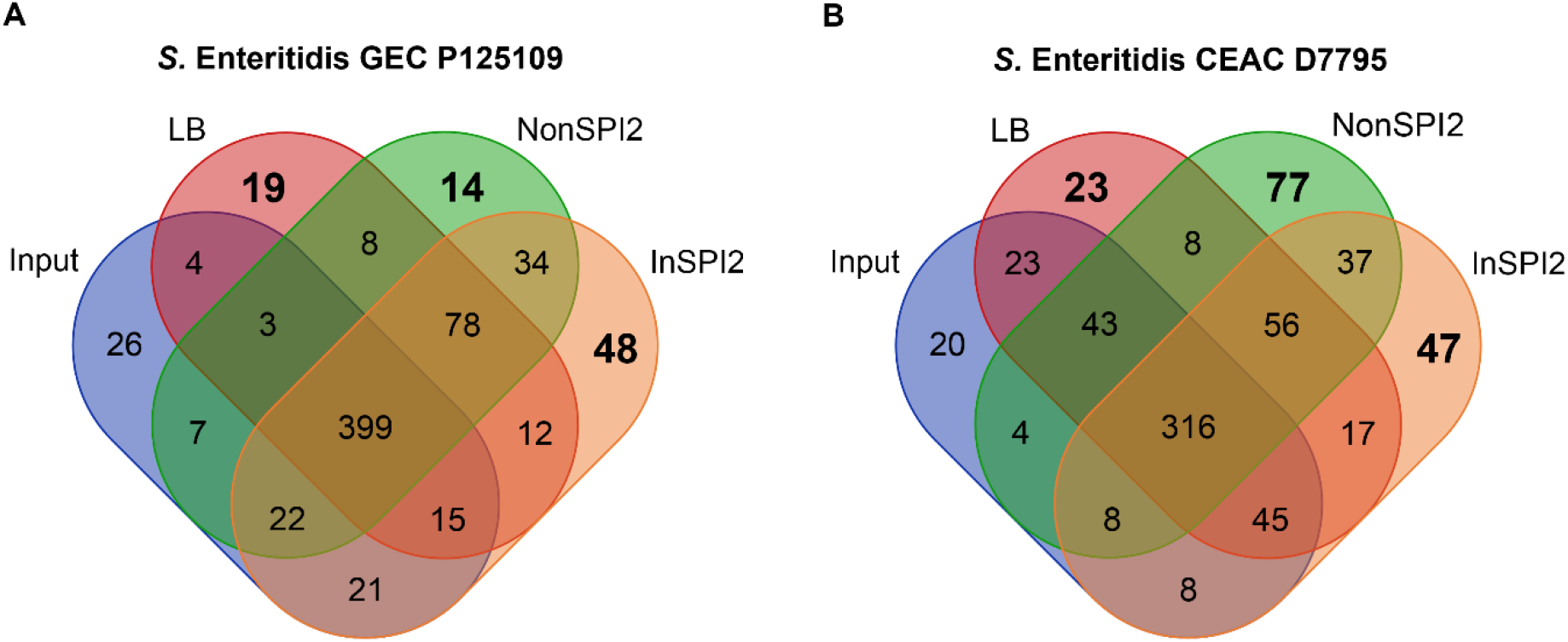
*S*. Enteritidis GEC P125109 and CEAC D7795 genes required for fitness in LB, NonSPI2 and InSPI2 media. The Venn diagrams compare the required genes in (A) P125109 or (B) D7795 with genes required for optimal growth in LB, NonSPI2 and InSPI2 media in the respective strains (Table S4). Generated using http://bioinformatics.psb.ugent.be/webtools/Venn/.

The P125109 LB-only required genes fall mainly into the major category of “Information Storage and Processing”, with 7 out of 19 genes (36%) belonging to this category, followed by 5 genes in the “Metabolism” major category (Table S4). Compared to the NonSPI2-only and InSPI2-only fitness genes, fewer metabolism-related genes were required for optimal growth in LB, which reflects the nutrient-rich LB environment [59]. Among the 19 LB-only required genes, 6 genes (*xseB*, *cydA*, *recB*, *gidA*, *yjeA*, *arcA*) have been previously identified as required by *S*. Typhi for growth in LB [26]. The *recB* gene, encoding an exonuclease subunit, was also reported to be required by *S*. Typhimurium 14028 [30, 60] and *E. coli* [27] after several passages in LB. The identification of *recB* gene in multiple TIS screens highlights the importance of this gene for fitness in the LB environment.

The P125109 genes that were required for optimal growth in NonSPI2 and/or InSPI2 were mostly metabolism-related genes, confirming the nutritional deficiencies encountered by the bacteria grown in synthetic minimal media. Genes required for optimal growth in both NonSPI2 and InSPI2 included *pur* genes (for purine biosynthesis) and *aro* genes (for aromatic amino acid biosynthesis). The InSPI2-only required genes included several genes associated with SPIs and RODs; the *dksA* gene was also identified, which plays a key role in the stringent response and has been experimentally validated as required for growth of *S*. Typhimurium in minimal medium [61, 62].

For CEAC strain D7795, 541 genes, 559 genes and 532 genes were designated as required in LB, NonSPI2 and InSPI2 following two passages in the respective growth media. After removing genes that had been identified as required in the input, there were 23 LB-only required genes, 77 NonSPI2-only required genes and 47 InSPI2-only required genes (Fig 3). As seen in P125109, most of the D7795 LB-only required genes were either involved in “Metabolism” (13 out 23 genes; 54%) or “Information Storage and Processing” (6 out of 23 genes; 25%) processes. Five genes identified as required in D7795 for LB growth had been reported to be required by *S*. Typhimurium previously: *ppiB*, a peptidyl-prolyl isomerase; *cydA*, cytochrome oxidase d subunit and *crp*, cAMP-activated global transcriptional regulator [63] and *rstB*, the sensor kinase of the *rstAB* two-component system, and *rnfD*, a component of the electron transport chain [31].

Most of the D7795 genes identified as required for optimal growth in NonSPI2 and InSPI2 minimal media were, as expected, metabolism-related, including genes involved in amino acid transport and metabolism (E), nucleotide transport and metabolism (F) and energy production and conversion (C) (Table S4). Similar to P125109, the D7795 *dksA* gene was also identified as required for growth in InSPI2 media. Several SPI-1 and SPI-2 genes and a SPI-1 effector SopE were required for growth in NonSPI2, while two SPI-5 genes (*pipB* and *sORF26*) and one ROD9 gene (*SEN1001*) were required for growth in InSPI2.

There were 78 and 56 genes designated as required in all growth media (LB, NonSPI2 and InSPI2) for P125109 and D7795, respectively (Fig 3 and Table S4). These genes represent the biological processes required by P125109 and D7795 for growth in laboratory conditions, and include genes such as *ftsK*, *icdA*, *rpiA*, *yheN*, *rfa* and *atp* genes, which are also required by *S*. Typhimurium ST19 14028 during *in vitro* growth [60, 64].

Overall, we conclude that similar functional categories of genes were required by both P125109 and D7795 for optimal growth in LB, NonSPI2 and InSPI2, respectively. These genes were mainly involved in energy production and conversion (C), carbohydrate transport and metabolism (G), amino acid transport and metabolism (E), nucleotide transport and metabolism (F) and transcription (K) (Fig S2).

### *S*. Enteritidis D7795 survives and replicates better in macrophages than *S*. Enteritidis P125109

To date, virulence phenotypes of CEAC D7795 have only been assessed in an avian infection model [9]. Survival and replication within macrophages is an important step in systemic *Salmonella* infections [24]. We compared the interaction of *S*. Enteritidis GEC P125109 and CEAC D7795 with RAW 264.7 macrophages, together with two additional *S*. Enteritidis strains, A1636 and CP255. A1636 is a GEC isolate from Africa, while CP255 belongs to the same Central/Eastern African clade as D7795 that originated from the Democratic Republic of Congo [9, 15]. Δ*phoPQ* deletion mutants were constructed in P125109 and D7795 by λ Red recombineering as negative controls for the infection studies. The PhoPQ two-component regulatory system is required for the survival of *S*. Typhimurium within macrophages [65, 66], and a previous genome-wide screen of *S*. Enteritidis P125109 Tn*5* mutants showed that *phoPQ* insertion mutants were negatively selected during murine infection [16]. The *S*. Typhimurium ST19 strains 4/74 [67] and 4/74 Δ*phoPQ* [68] were also included for comparison. All *S*. Enteritidis and *S*. Typhimurium strains tested in this experiment were taken up by RAW 264.7 macrophages at similar levels. Importantly, we found that the CEAC isolates showed significantly higher levels of intra-macrophage replication than GEC isolates at 15.5 h post-infection (Fig 4).

**Fig 4.**
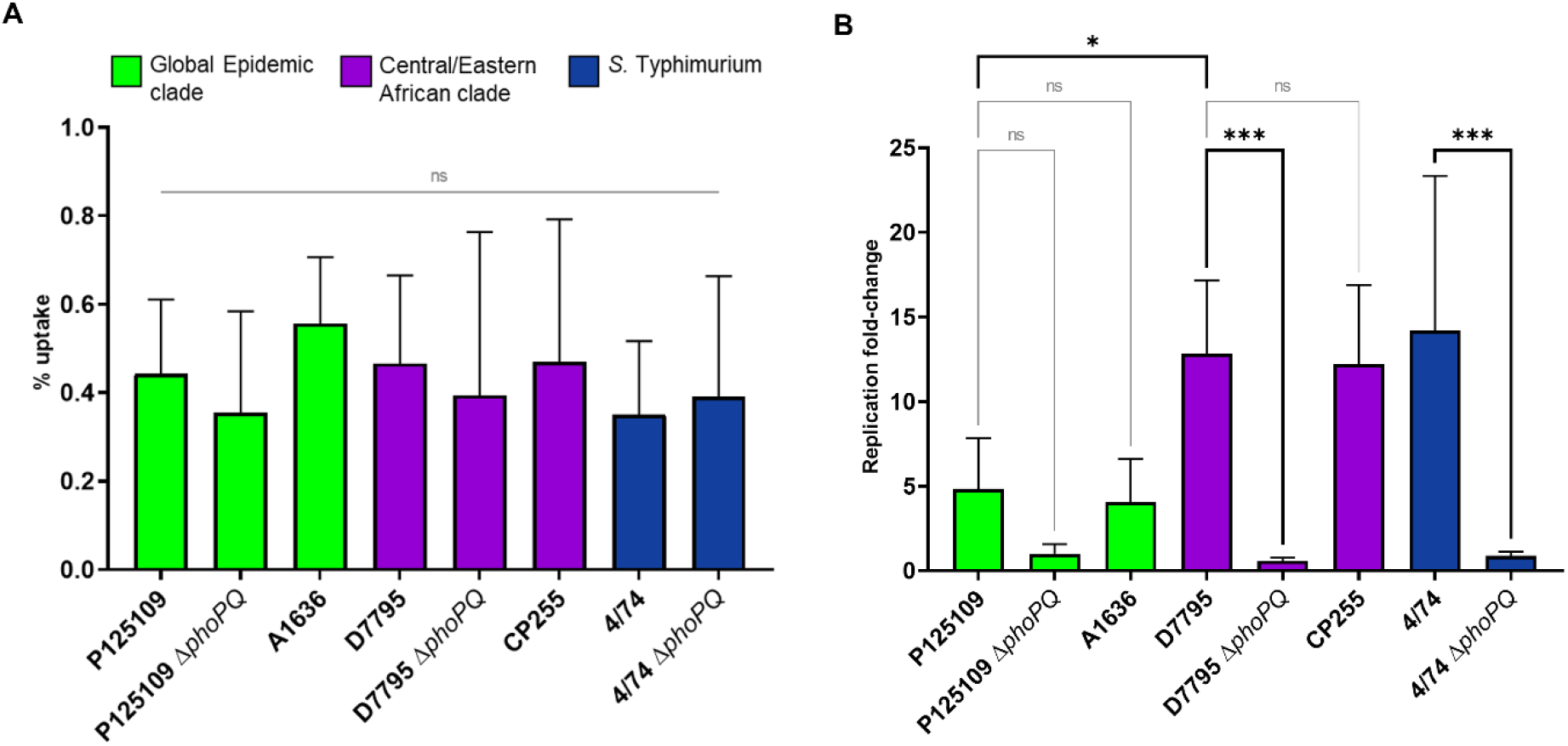
*S*. Enteritidis CEAC D7795 displays higher levels of intra-macrophage survival and replication than *S*. Enteritidis GEC P125109. *S*. Enteritidis strains are colour-coded by clades, as defined by Feasey *et al*. [9]: green, GEC; purple, CEAC. The *S*. Typhimurium 4/74 and 4/74 Δ*phoPQ* (blue) strains were included as additional controls. (A) Uptake of *Salmonella* strains by RAW 264.7 macrophages, shown as the percentage of the infecting inoculum recovered (CFU) at 1.5 h post-infection (p.i.). (B) Intra-macrophage replication of *Salmonella*, shown as the ratio of intracellular bacteria (CFU) recovered at 15.5 h p.i. compared to the CFU recovered at 1.5 h p.i., as fold-change. Both panels (A and B) represent average values obtained from five independent experiments with three replicates each, and error bars show standard deviation. Statistical tests used were one-way ANOVA, followed by Bonferroni’s multiple comparison test to compare selected pairs of means. ns, *P* ≥ 0.05; *, *P* < 0.05; ***, *P* < 0.001.

### Identification of fitness genes of *S*. Enteritidis P125109 and D7795 following macrophage infection

Having established that the CEAC strain D7795 survives and replicates better in macrophages than GEC strain P125109, the respective transposon libraries were used to investigate the process of intracellular infection of RAW 264.7 macrophages. Each pool of transposon mutants was grown in LB (Input_LB) then passaged once through murine macrophages. Intra-macrophage bacteria were recovered at 12 h post-infection and grown in LB for 10 h to generate the output culture (Output_MAC). A fraction of the input was sub-cultured in LB for 10 h to ascertain the effect of growth in LB broth culture on the composition of the transposon library (Output_LB_10h). Genomic DNA from the input and output samples was purified and prepared for Illumina sequencing of DNA adjacent to the transposon, as described in Materials and Methods. A total of 12 DNA libraries were sequenced on a single lane of an Illumina HiSeq4000, generating a total of 340,375,208 paired-end reads. The number of sequence reads that contained the transposon tag sequence, and the sequence reads that were uniquely mapped to the respective *S*. Enteritidis genomes are presented in Table S2.

Genes that modulated the intracellular survival and replication of *Salmonella* in RAW 264.7 macrophages were identified by comparing the macrophage output samples with the input samples and calculating the changes in frequency of reads mapped to each gene, expressed as log_2_(fold change) (FC) [31]. A gene is considered to exhibit differential fitness if its log_2_FC value is less than 1 (attenuated fitness) or greater than 1 (increased fitness) <u>with</u> a *P*-value < 0.05. Genes affecting growth in LB for 10 h were similarly identified by comparing the LB output samples with the input samples. Required genes were identified from the input sample using the insertion index and excluded from the list of differential fitness genes.

Following 12-h macrophage infection, a total of 479 P125109 genes were identified as required from essentiality analysis of the input libraries. In total, transposon insertions in 327 genes were associated with attenuated replication within macrophages (Fig 5 and Table S5). To identify genes important for fitness inside macrophages but not for growth in laboratory media, we first compared the 327 macrophage-attenuated genes with the 124 genes that showed attenuation in 10-h LB growth when disrupted by a transposon insertion (Table S5). This analysis identified 227 genes that only attenuated fitness of P125109 during macrophage infection, and not during 10-h growth in LB (Fig S3 and Table S4).

**Fig 5.**
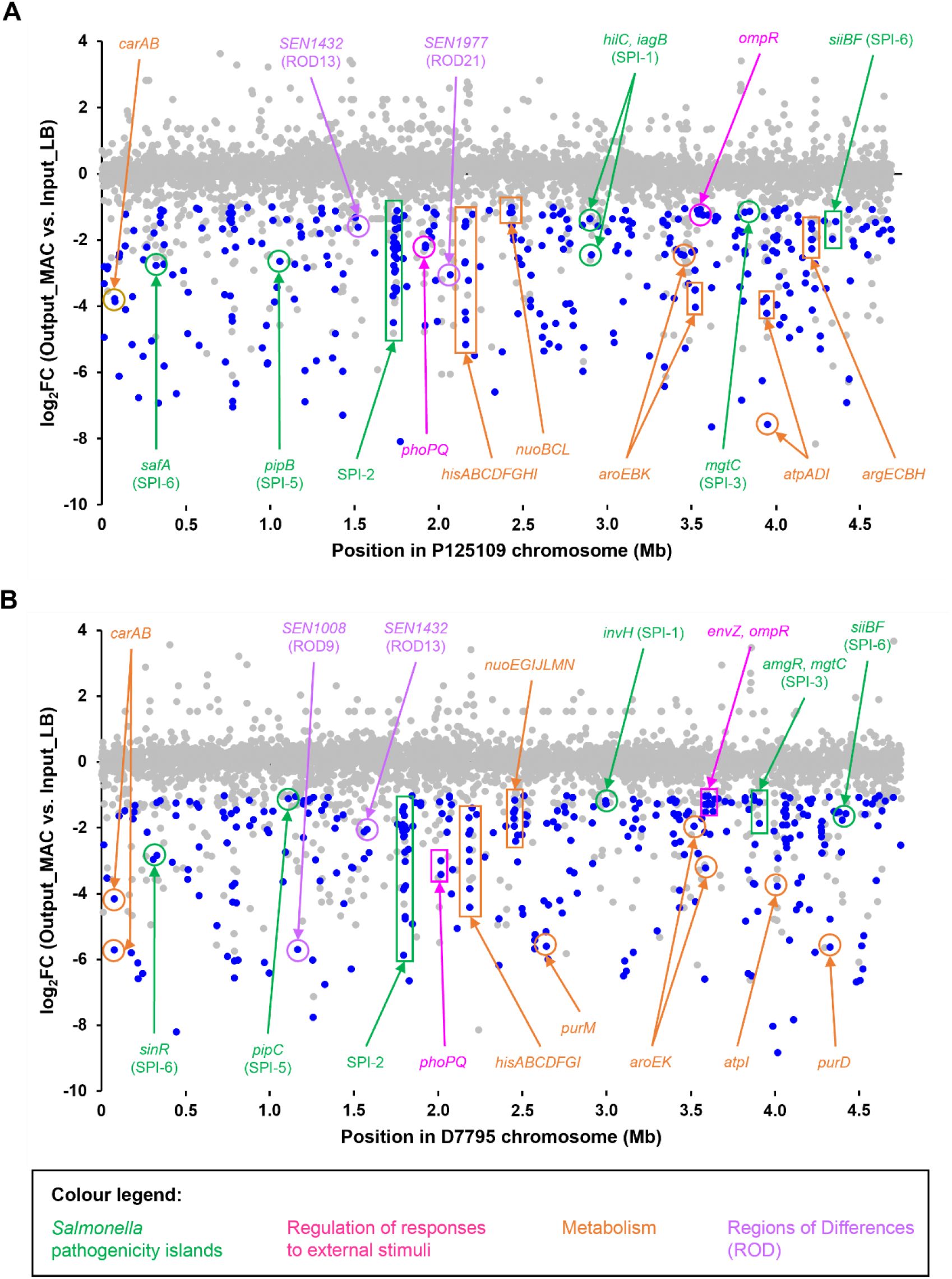
Macrophage-attenuated genes of *S*. Enteritidis GEC P125109 and CEAC D7795. The *S*. Enteritidis GEC P125109 and CEAC D7795 transposon libraries were used to infect RAW 264.7 macrophages for 12 h. Relative abundance of each mutant was determined by comparing the frequency of sequenced reads mapped to each gene after the infection (Output_MAC) to the initial inoculum (Input_LB). Each position on the *x*-axis represents the starting nucleotide position of each gene locus on the *Salmonella* chromosome, and the *y-*axis represents the log_2_(fold change) of changes in abundance of mapped reads. Loci with significant reduction in abundance (log_2_FC < −1, *P* < 0.05) are shown in blue (attenuated fitness). Grey dots include both loci with significant increase in abundance (log_2_FC > 1, *P* < 0.05) and loci with *P* ≥ 0.05. DeSeq2 analysis of the TIS macrophage data is presented in Table S5.

The resulting 227 genes were then cross-referenced with genes required for growth in the LB, NonSPI2 and InSPI2 *in vitro* laboratory conditions tested in this study (Fig S3). We identified a total of 320 genes that were “macrophage-associated”. Of these, 177 genes were “macrophage-specific”, only having reduced fitness during macrophage infection with no impact upon growth *in vitro* (Fig S3 and Table S4). The terms “macrophage-specific” and “macrophage-associated” have been defined previously [31], and are explained in the legend to Fig S3.

The 177 “macrophage-specific” P125109 genes included many well-characterised genes important for intra-macrophage survival of *S*. Typhimurium, such as 22 SPI-2 genes, *mgtC* from SPI-3, and several global regulatory systems that control *Salmonella* virulence (e.g. *ompR*, *phoQ*, *ssrB*). Two SPI-1 genes, *hilC* and *iagB*, were also identified. There were 74 genes related to various metabolic processes, including arginine biosynthesis (*arg* genes), histidine biosynthesis (*his*) genes, the TCA cycle (*sucD*) and oxidative phosphorylation (*ndh*), reflecting the nutritional stresses encountered by the bacteria within the macrophage environment. Three genes only present in P125109 were also identified as “macrophage-specific”, namely *SEN0912* (encoding a hypothetical protein) and two tRNA genes (*tRNA-Ala* and *tRNA-Thr*). No genes from the ROD9 region were identified, despite reports that ROD9-associated genes played a role in virulence in macrophage and animal infection [16, 20, 69]. Based on COG classifications, the majority of the 177 “macrophage-specific” genes were associated with nucleotide transport and metabolism (F, 17%), amino acid transport and metabolism (E, 10%), transcription (K, 8%) and translation (J, 9%).

Genes important for intra-macrophage fitness of D7795 were identified as described earlier. Essentiality analysis of the input sample identified 432 required genes, and these genes were excluded from the list of genes that caused differential fitness in macrophage infection. Transposon insertions in 329 genes caused attenuation (log_2_FC < −1, *P* < 0.05) during RAW macrophage infection (Fig 5). Of the 329 genes, 78 genes exhibited reduced fitness during growth in LB for 10 h. Cross-referencing the 251 genes with the genes required for *in vitro* growth under laboratory conditions identified a total of 325 “macrophage-associated” genes and 201 “macrophage-specific” genes (Fig S3 and Table S4). Similar to the findings for P125109, the 201 D7795 “macrophage-specific” genes are primarily located in the SPI regions. Specifically, most of the attenuated mutants were located in SPI-2; several others were identified in SPI-3 (*SEN3578*, *amgR*, *mgtC*), SPI-4 (*siiF*), SPI-5 (*pipC*), SPI-6 (*SEN0286*) and SPI-10 (*SEN4249*). Transposon insertions in *SEN1008* from ROD9 (SPI-19) also attenuated macrophage fitness. Distribution of the COG functional categories of the “macrophage-specific” genes is broadly similar to that in P125109, with nucleotide transport and metabolism (F) and amino acid transport and metabolism (E) genes forming the largest categories.

Transposon insertion in 61 P125109 and 18 D7795 genes resulted in increased fitness in macrophage infection (log_2_FC > 1, *P* < 0.05), and include the *rfa*/*rfb* genes that are responsible for lipopolysaccharide (LPS) O-antigen biosynthesis (Table S5). Increased intra-macrophage fitness conferred by transposon disruption in the *rfa*/*rfb* genes was also observed in *S*. Typhimurium infection of RAW 264.7 macrophages [31, 70]. The *rfb* genes are involved in O-antigen synthesis while the *rfa* genes mediate LPS core synthesis [71]. As noted by Canals *et al*. [31], *Salmonella* mutants lacking the LPS O-antigen are phagocytosed at higher levels than wild-type strains by murine macrophages [72]. This likely accounts for the increased number of *S*. Enteritidis *rfa*/*rfb* mutants recovered in the macrophage output, and reiterates the importance of interpreting the results of a mutant screen in the context of the experimental model used [63, 70].

### *S*. Enteritidis P125109 and D7795 genes that modulate intra-macrophage fitness are required for virulence in other *Salmonella* serovars

The *S*. Enteritidis GEC P125109 and CEAC D7795 genes associated with differential fitness in macrophage infection were compared with other infection models and *Salmonella* serovars. Genes affecting intra-macrophage fitness of *S*. Typhimurium ST313 D23580 [31] were identified by re-processing and analysis of the data using pipelines described in Materials and Methods. Genes required for *Salmonella* infection in other serovars and/or infection models were retrieved from published data sets using parameters established by the authors of the original papers [16, 70, 73, 74]. Only orthologous genes were used in the comparisons.

We found that mutation of a total of 147 genes caused attenuated macrophage infection for *S*. Enteritidis P125109, D7795 and *S*. Typhimurium ST313 D23580 (Fig 6 and Table S6). There was also significant overlap with genes required for *Salmonella* virulence in mice and other infection models, demonstrating that *S*. Enteritidis shares many virulence genes in common with other serovars (Fig 6). Many of these genes include the well-characterised regulatory systems that control *Salmonella* virulence (the *phoPQ* two-component regulators and *ompR*, an element of the *ompR-envZ* two-component regulatory system), and SPI-2 genes that encode structural components of the type III secretion system, and genes involved in purine and aromatic amino acids biosynthesis (Table S6 and Fig S4).

**Fig 6.**
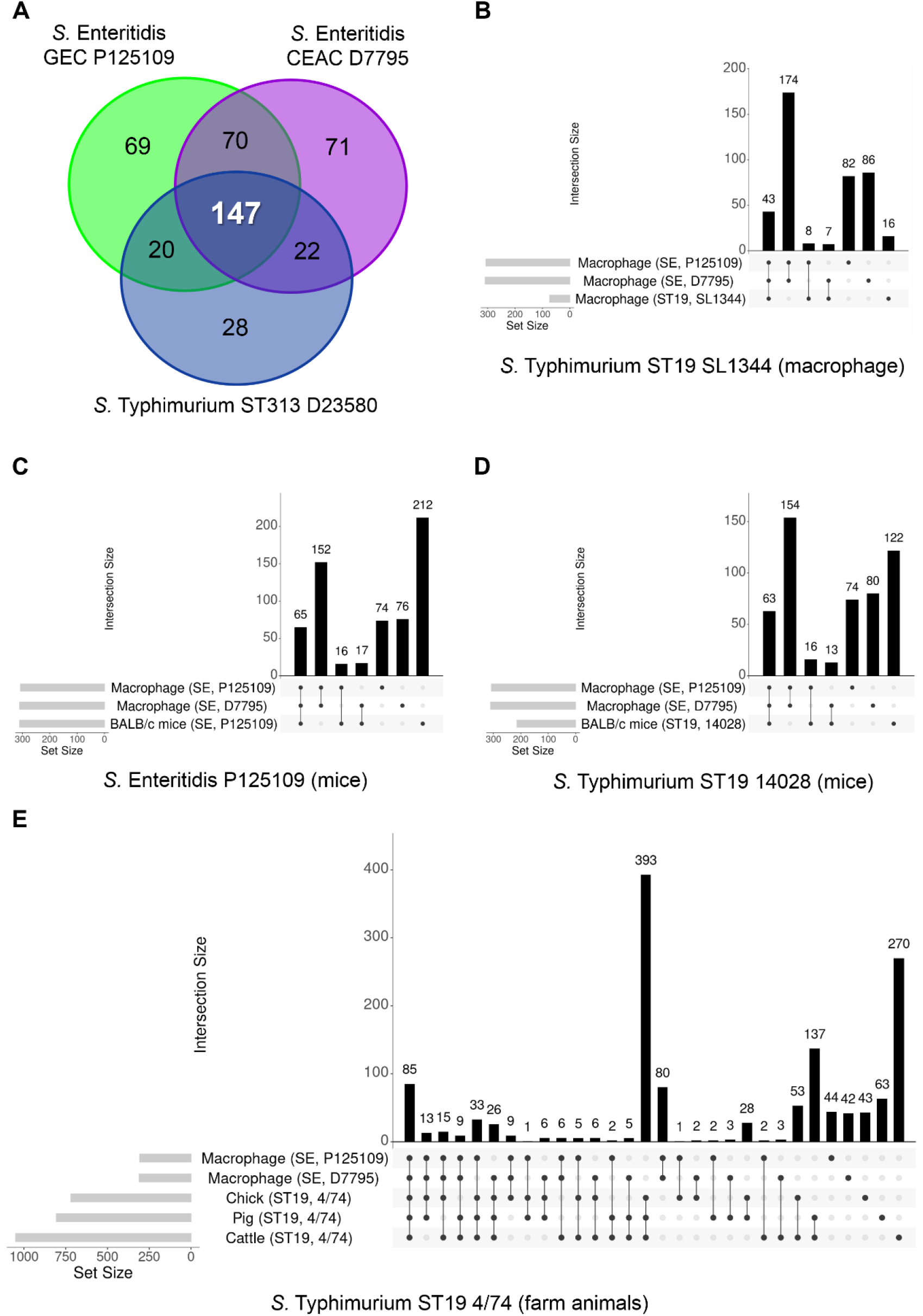
Genes that modulate intra-macrophage fitness of *S*. Enteritidis GEC P125109 and CEAC D7795 are required for virulence in other infection models. Macrophage-attenuated genes in *S*. Enteritidis GEC P125109 and CEAC D7795 were compared with genes associated with virulence in macrophage for (A) *S*. Typhimurium ST313 D23580 [31]; (B) ST19 SL1344 [70]; P125109 [16] and for *S*. Typhimurium ST19 14028 in BALB/c mice [73]; and for *S*. Typhimurium ST19 4/74 in food-related animal infection models [74]. Only orthologous genes were included in the analyses (Table S6). UpSet plots were generated using Intervene [75].

### *S*. Enteritidis D7795 and *S*. Typhimurium ST313 D23580 do not share novel virulence factors

We used the TIS data to search for genes involved in intra-macrophage virulence that were only found in African *Salmonella* strains. Focusing on the three-way comparison between *S*. Enteritidis GEC P125109, CEAC D7795 and *S*. Typhimurium ST313 D23580, 22 genes were identified as modulating macrophage fitness in invasive African *Salmonella* strains D7795 and D23580 but not in gastroenteritis-associated P125109 (Fig 6). Cross-referencing these 22 genes with *Salmonella* virulence genes identified previously [16, 70, 73, 74] showed that all 22 genes have been implicated in at least one other infection model (Table S6). We conclude that no novel virulence factors that modulate intra-macrophage fitness are shared by the two African *Salmonella* strains.

### Candidate novel macrophage fitness genes that were unique to *S*. Enteritidis P125109 or D7795

The three-way comparison between *S*. Enteritidis GEC P125109, CEAC D7795 and *S*. Typhimurium ST313 D23580 identified mutations in 69 P125109 genes and 71 D7795 genes that attenuated intra-macrophage fitness (Fig 6). We investigated the role of these 69 and 71 genes in other infection models or in *S*. Typhimurium, and did a detailed comparison against genes identified in other published studies [16, 70, 73, 74]. We identified a total of 22 P125109 genes and 39 D7795 genes that were only associated with attenuated macrophage fitness in a single strain (Tables 1 and 2). These 22 and 39 genes represent candidate novel virulence genes for P125109 and D7795 respectively, and include strain-specific genes (i.e. genes without orthologues in the other *Salmonella* strains referenced in this study) (e.g. *SEN0912* in P125109 and three tRNA genes of D7795, locus tags D7795_02738, D7795_03122 and D7795_04774) and genes with orthologues in the other *Salmonella* strains and serovars that were not associated with attenuated fitness during macrophage infection (e.g. *lipB*∷Tn*5*, *gatR*∷Tn*5*, *hscB*∷Tn*5*, *rnfE*∷Tn*5* and *sopD2*∷Tn*5* are attenuated in D7795 but not in P125109, *S*. Typhimurium D23580 and/or LT2). Experimental validation of these candidates by individual gene deletion mutation will be necessary to verify a role in intra-macrophage replication.

**Table 1.** Candidate novel macrophage fitness genes in *S*. Enteritidis P125109.

| Name | Product | P125109 ID | SEN ID | D7795 ID | D23580 ID | LT2 ID | P125109 log<sub>2</sub>FC |
| --- | --- | --- | --- | --- | --- | --- | --- |
| No orthologue in D7795, D23580 or LT2 |  |  |  |  |  |  |  |
| *SEN0912* | hypothetical protein | P125109_01026 | SEN0912 | NONE | NONE | NONE | -1.41 |
| *tRNA-Ala* | tRNA-Ala(ggc) | P125109_02722 | #N/A | #N/A | #N/A | #N/A | -2.28 |
| *tRNA-Thr* | tRNA-Thr(ggt) | P125109_04470 | #N/A | #N/A | #N/A | #N/A | -3.47 |
| Present in D7795, D23580 and/or LT2 but only attenuated in P125109 |  |  |  |  |  |  |  |
| *tpke11* | NONE | P125109_00013 | NONE | D7795_00014 | STMMW_ncRNA_1 | NONE | -3.32 |
| *yabC* | conserved hypothetical protein | P125109_00141 | SEN0121 | D7795_00143 | STMMW_01261 | STM0120 | -1.23 |
| *yaeD* | conserved hypothetical protein | P125109_00284 | SEN0256 | D7795_00285 | STMMW_02532 | STM0248 | -5.05 |
| *SEN0328* | hypothetical protein | P125109_00373 | SEN0328 | D7795_00374 | STMMW_04151 | STM0345 | -2.14 |
| *SEN0532* | hypothetical protein | P125109_00595 | SEN0532 | D7795_00602 | STMMW_06191 | STM0551 | -1.64 |
| *sucD* | succinyl-CoA synthetase alpha chain | P125109_00774 | SEN0689 | D7795_00799 | STMMW_07961 | STM0739 | -1.35 |
| *SEN0907* | putative ion:amino acid symporter | P125109_01022 | SEN0907 | D7795_01047 | STMMW_10141 | STM1003 | -1.01 |
| *yedD* | putative lipoprotein | P125109_01175 | SEN1044 | D7795_01257 | STMMW_19441 | STM1964 | -3.78 |
| *cheZ* | chemotaxis protein CheZ | P125109_01224 | SEN1088 | D7795_01307 | STMMW_18981 | STM1915 | -1.81 |
| *lrhA* | NADH dehydrogenase operon transcriptional regulator | P125109_02623 | SEN2312 | D7795_02672 | STMMW_23521 | STM2330 | -1.57 |
| *iagB* | cell invasion protein | P125109_03091 | SEN2719 | D7795_03189 | STMMW_28391 | STM2877 | -2.45 |
| *ygbK* | conserved hypothetical protein | P125109_03136 | SEN2756 | D7795_03234 | STMMW_28801 | STM2917 | -1.59 |
| *sORF75* | NONE | P125109_03192 | NONE | D7795_03290 | STMMW_29261 | NONE | -1.75 |
| *yggT* | putative membrane protein | P125109_03353 | SEN2944 | D7795_03449 | STMMW_30621 | STM3101 | -1.50 |
| *glpR* | glycerol-3-phosphate regulon repressor | P125109_03805 | SEN3348 | D7795_03904 | STMMW_35121 | STM3523 | -1.25 |
| *prlC* | oligopeptidase A | P125109_03882 | SEN3417 | D7795_03981 | STMMW_35831 | STM3594 | -1.28 |
| *wecG* | probable UDP-N-acetyl-D-mannosaminuronic acid transferase | P125109_04235 | SEN3734 | D7795_04333 | STMMW_39041 | STM3929 | -1.26 |
| *serB* | putative phosphoserine phosphatase | P125109_04914 | SEN4334 | D7795_05011 | STMMW_45211 | STM4578 | -1.38 |
| *yjjY* | conserved hypothetical protein | P125109_04937 | SEN4355 | D7795_05034 | STMMW_45421 | STM4599 | -2.03 |

P125109 ID = P125109 locus tags from re-annotation [15]; SEN ID = *S. Enteritidis* identifiers;D23580 = *S. Typhimurium* ST313 strain D23580; LT2 = *S. Typhimurium* ST19 strain LT2

**Table 2.**
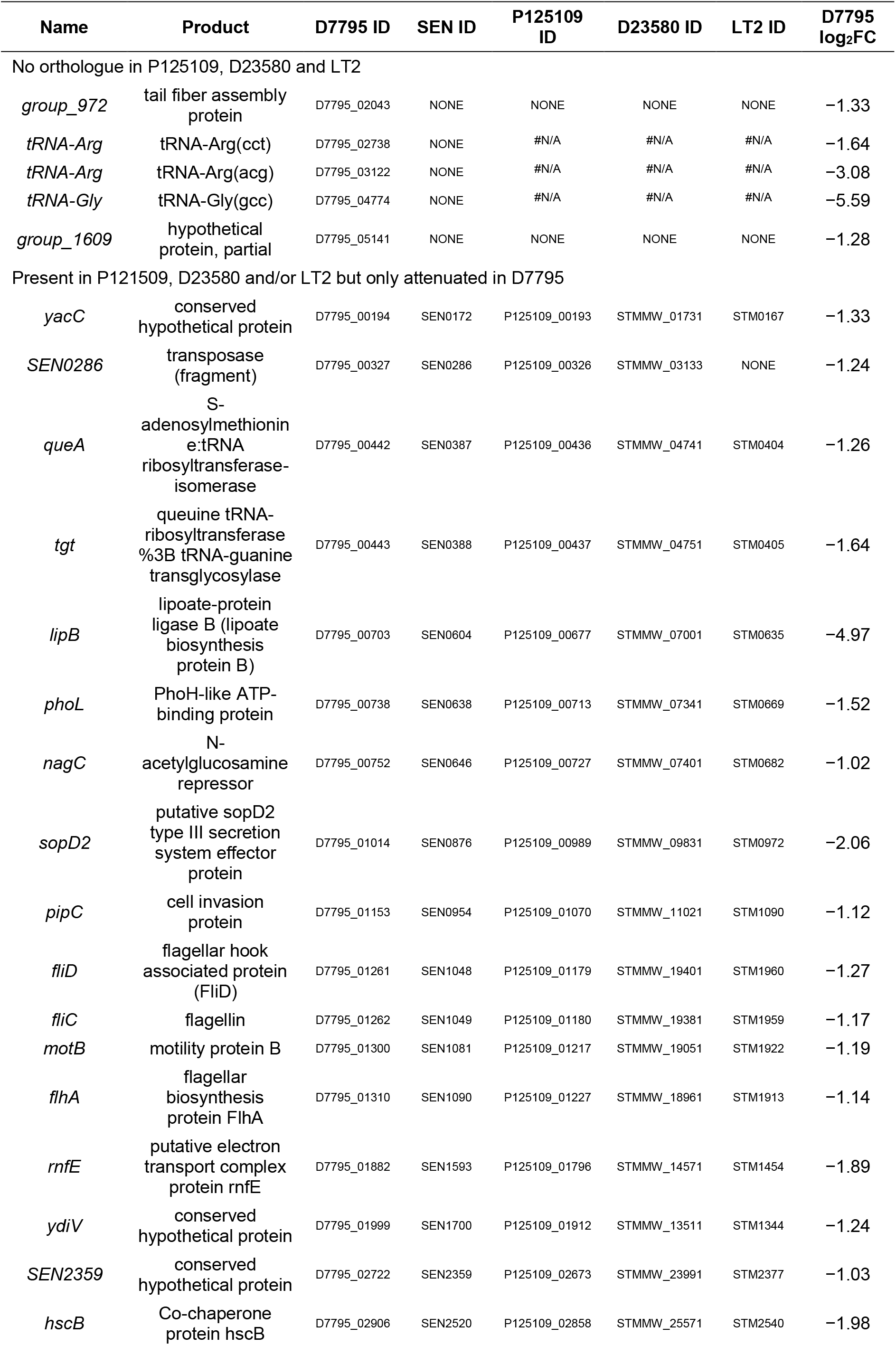
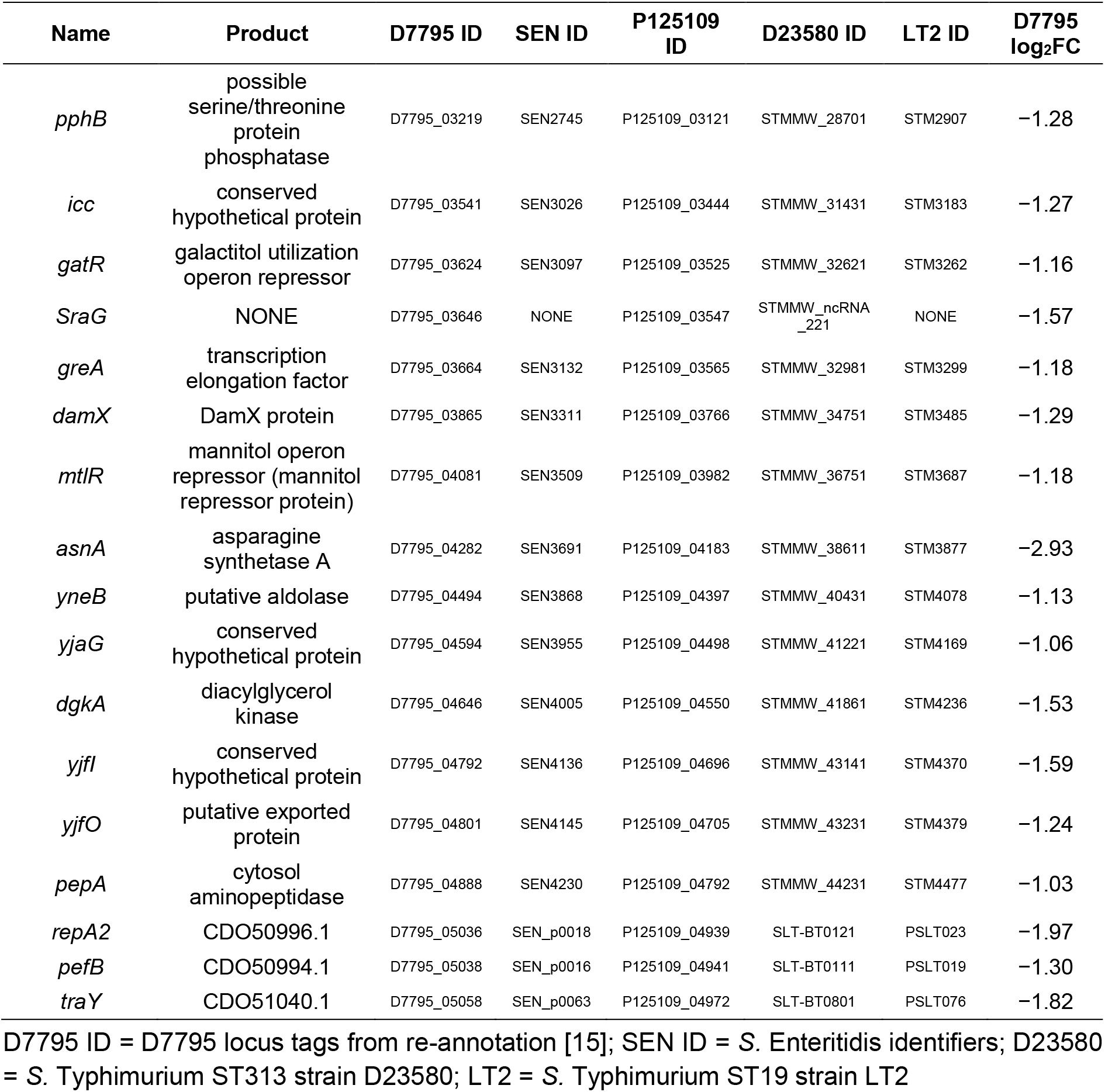
Candidate novel macrophage fitness genes in *S*. Enteritidis D7795.

## PERSPECTIVE

Since the identification of genetic variants of *S*. Typhimurium and *S*. Enteritidis that are highly associated with bloodstream infections in sub-Saharan Africa, only the *S*. Typhimurium pathovariant has been an active focus of research [76]. Functional genomic studies involving African *S*. Enteritidis have lagged behind. Here, we used a global mutagenesis approach to compare gene function between bloodstream infection-associated *S*. Enteritidis CEAC strain D7795 and gastroenteritis-associated GEC strain P125109. Our findings reveal broad similarities between the gene sets required for growth under laboratory conditions and macrophage infection by P125109 and D7795. The majority of these genes were also important for fitness in other *Salmonella* serovars and infection models. We identified 39 genes that could encode candidate novel virulence factors for *S*. Enteritidis CEAC strain D7795 and are worthy of further investigation.

## ACKNOWLEDGEMENTS

We are grateful to present and former members of the Hinton laboratory for valuable discussions, particularly Yan Li and Hermione Webster, and appreciated Paul Loughnane’s masterful technical assistance. We thank John Kenny, Pia Koldkjær, Anita Lucaci and Charlotte Nelson for their expertise in library preparation for Illumina sequencing, and the Centre for Genomic Research at the University of Liverpool for the use of the BioRuptor@Pico sonication system.

## AUTHOR CONTRIBUTIONS

Conceptualization: W.Y.F and R.C.; Data curation: W.Y.F, A.P. and B.P.S; Formal analysis: W.Y.F and A.P.; Funding acquisition: J.C.D.H.; Investigation: W.Y.F. and A.P.; Methodology: W.Y.F and R.C.; Project administration: W.Y.F. and J.C.D.H.; Resources: N.W., L.L., N.F. and P.W.; Software: A.P.; Supervision: W.Y.F., P.W. and J.C.D.H.; Validation: W.Y.F; Visualisation: W.Y.F and A.P.; Writing – original draft: W.Y.F and J.C.D.H.; Writing – review & editing: W.Y.F, R.C., A.P., B.P.S, N.W., L.L., N.F., P.W. and J.C.D.H.

## FUNDING INFORMATION

This work was supported by a Wellcome Trust Senior Investigator award [grant number 106914/Z/15/Z] to J.C.D.H. For the purpose of open access, the authors have applied a CC BY public copyright licence to any Author Accepted Manuscript version arising from this submission.

## CONFLICTS OF INTEREST

R.C. was employed by the University of Liverpool at the time of the study and is now an employee of the GSK group of companies.

## SUPPLEMENTARY DATA

**Fig S1.**
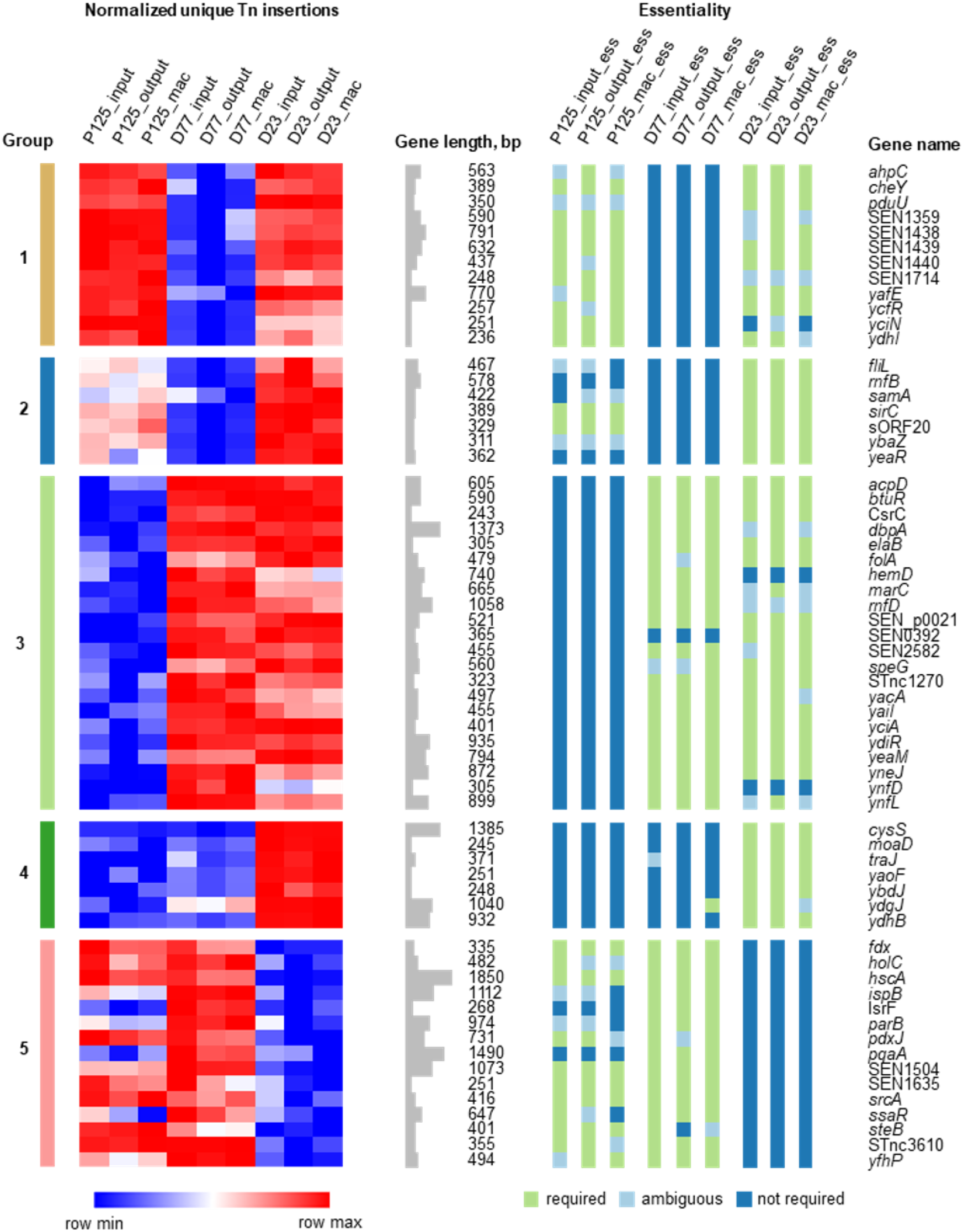
Inter-strain essentiality analysis identifies 63 genes that are differentially required between *S*. Enteritidis P125109, *S*. Enteritidis D7795 and *S*. Typhimurium D23580. Transposon insertion read counts are represented as a heat map, with red indicating many insertions and blue indicating very few insertions. Essentiality calls (based on insertion indices) from LB input, LB output and macrophage output libraries are represented by green and blue squares. Samples included: P125_input (P125109 LB input), P125109_output (P125109 LB output), P125_mac (P125109 macrophage output), D77_input (D7795 LB input), D77_output (D7795 LB output), D77 mac (D7795 macrophage output), D23_input (D23580 LB input), D23_output (D23580 LB output) and D23_mac (D23580 macrophage output).

**Fig S2.**
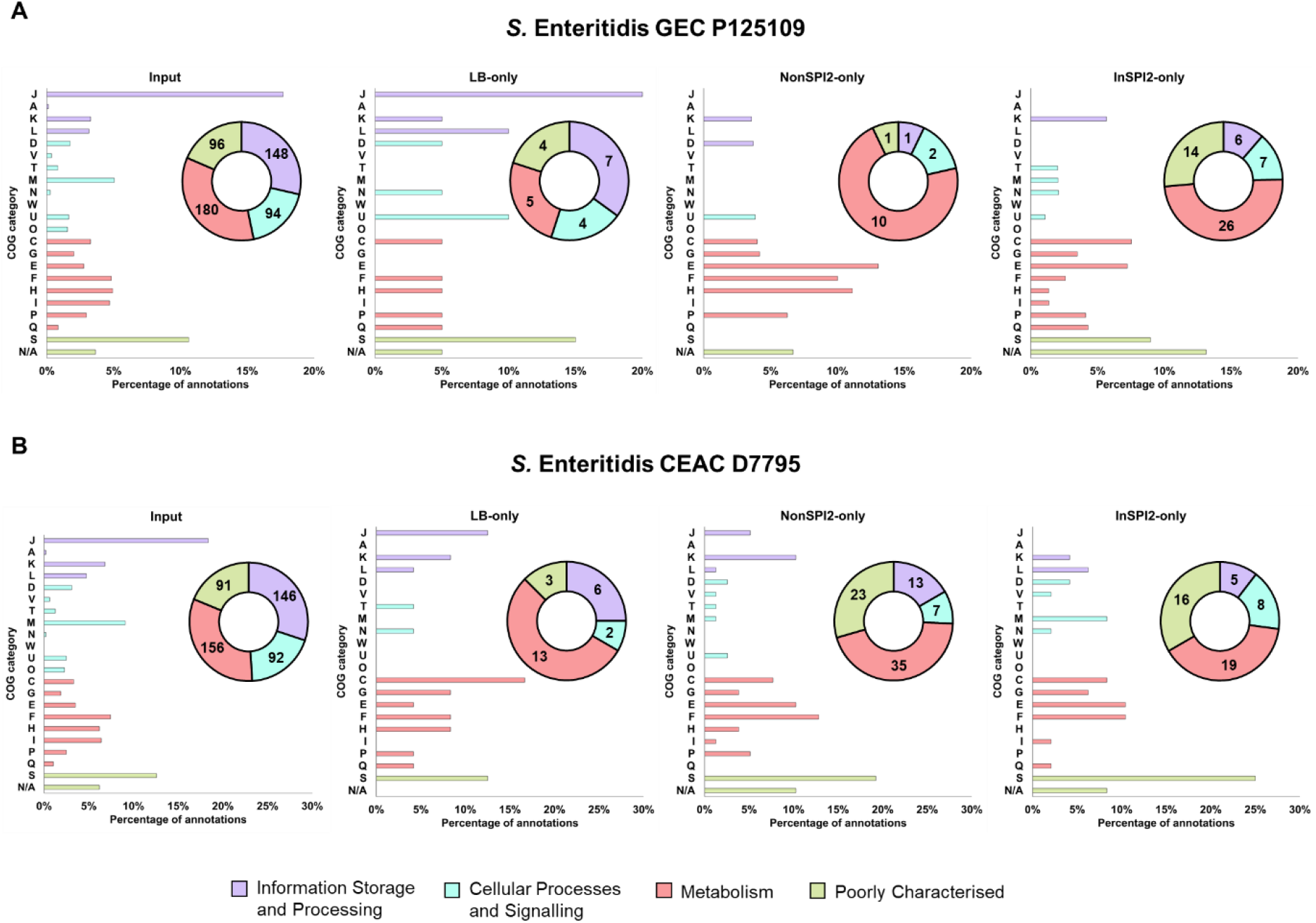
Distribution of Cluster of Orthologous Genes (COG) annotations in genes required by *S*. Enteritidis P125109 and D7795 for optimal growth in LB, NonSPI2 and InSPI2. Doughnut charts (insets) show the distribution of COG annotations in the four major functional categories (Information Storage and Processing, Cellular Processes and Signalling, Metabolism, Poorly Characterised), with total numerical counts in each major category shown. “Input” bar chart shows COG annotations from all genes identified as required in the respective *S*. Enteritidis Input libraries (497 genes for P125109 and 467 genes for D7795), whereas the “LB-only”, “NonSPI2-only” and “InSPI2-only” bar charts considered only the genes specific to that growth media (Fig 3). COG categories: J, Translation, ribosomal structure and biogenesis; A, RNA processing and modification; K, Transcription; L, Replication, recombination and repair; D, Cell cycle control, cell division, chromosome partitioning; V, Defense mechanisms; T, Signal transduction mechanisms; M, Cell wall/membrane/envelope biogenesis; N, Cell motility; W, Extracellular structures; U, Intracellular trafficking, secretion, and vesicular transport; O, Posttranslational modification, protein turnover, chaperones; C, Energy production and conversion; G, Carbohydrate transport and metabolism; E, Amino acid transport and metabolism; F, Nucleotide transport and metabolism; H, Coenzyme transport and metabolism, I, Lipid transport and metabolism; P, Inorganic ion transport and metabolism; Q, Secondary metabolites biosynthesis, transport and catabolism; S, Function unknown; N/A, not assigned.

**Fig S3.**
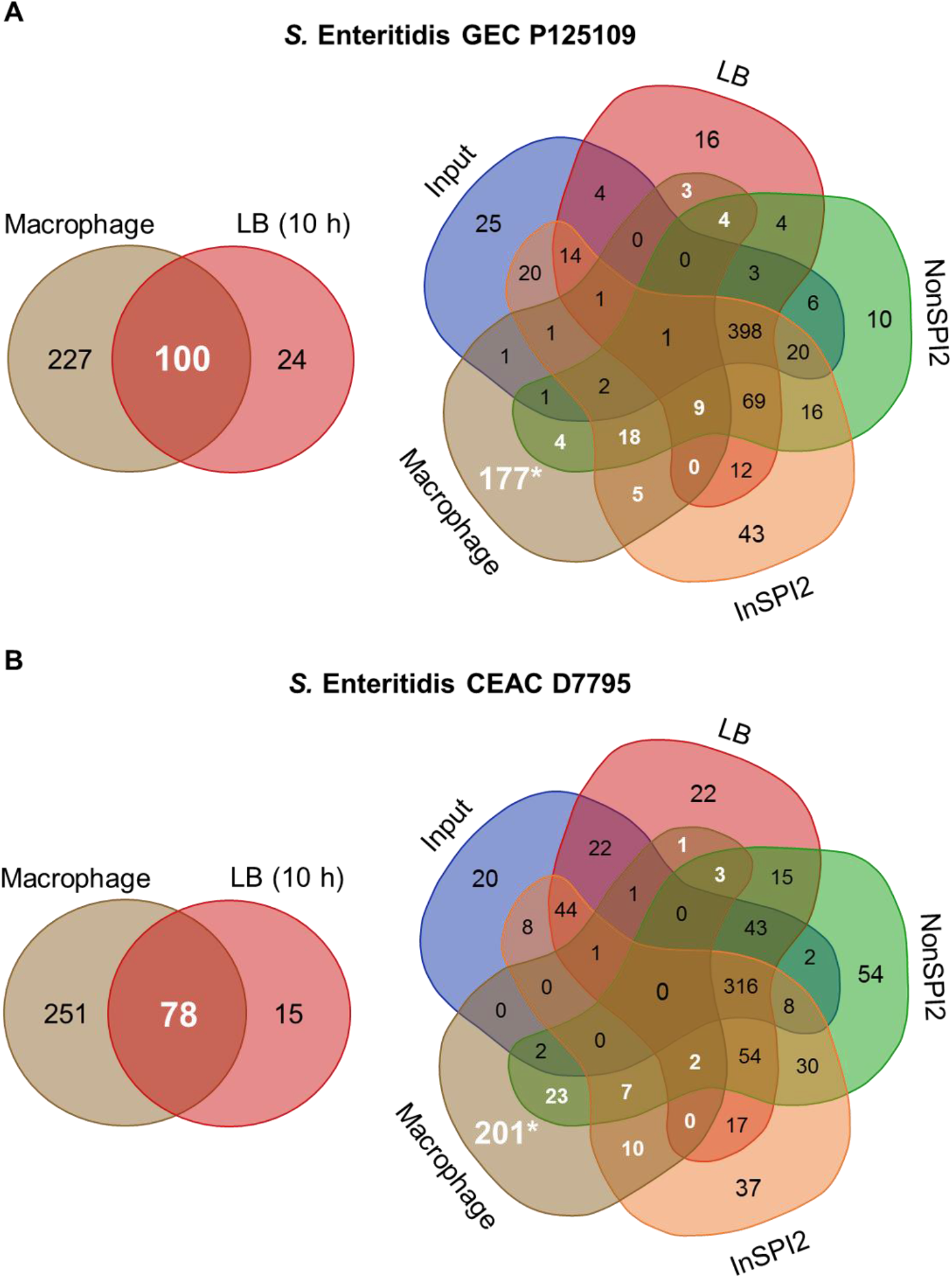
Macrophage-specific and macrophage-associated genes in *S*. Enteritidis P125109 and D7795. Genes that modulated the intracellular survival and replication of *Salmonella* in RAW 264.7 macrophages (identified by log_2_FC [Output_MAC vs. Input_LB] < −1 and *P*-value < 0.05) were compared with genes affecting growth in LB for 10 h (two-way Venn diagram in both panels A and B). The resulting 227 and 251 macrophage-attenuated fitness mutants in P125109 and D7795, respectively, were then compared with the genes required for *in vitro* growth under laboratory conditions (five-way Venn diagram). The numbers highlighted in white represent the “macrophage-associated” genes, and the numbers highlighted with an asterisk (*) represent the “macrophage-specific” genes. Venn diagrams were generated using http://bioinformatics.psb.ugent.be/webtools/Venn/.

**Fig S4.**
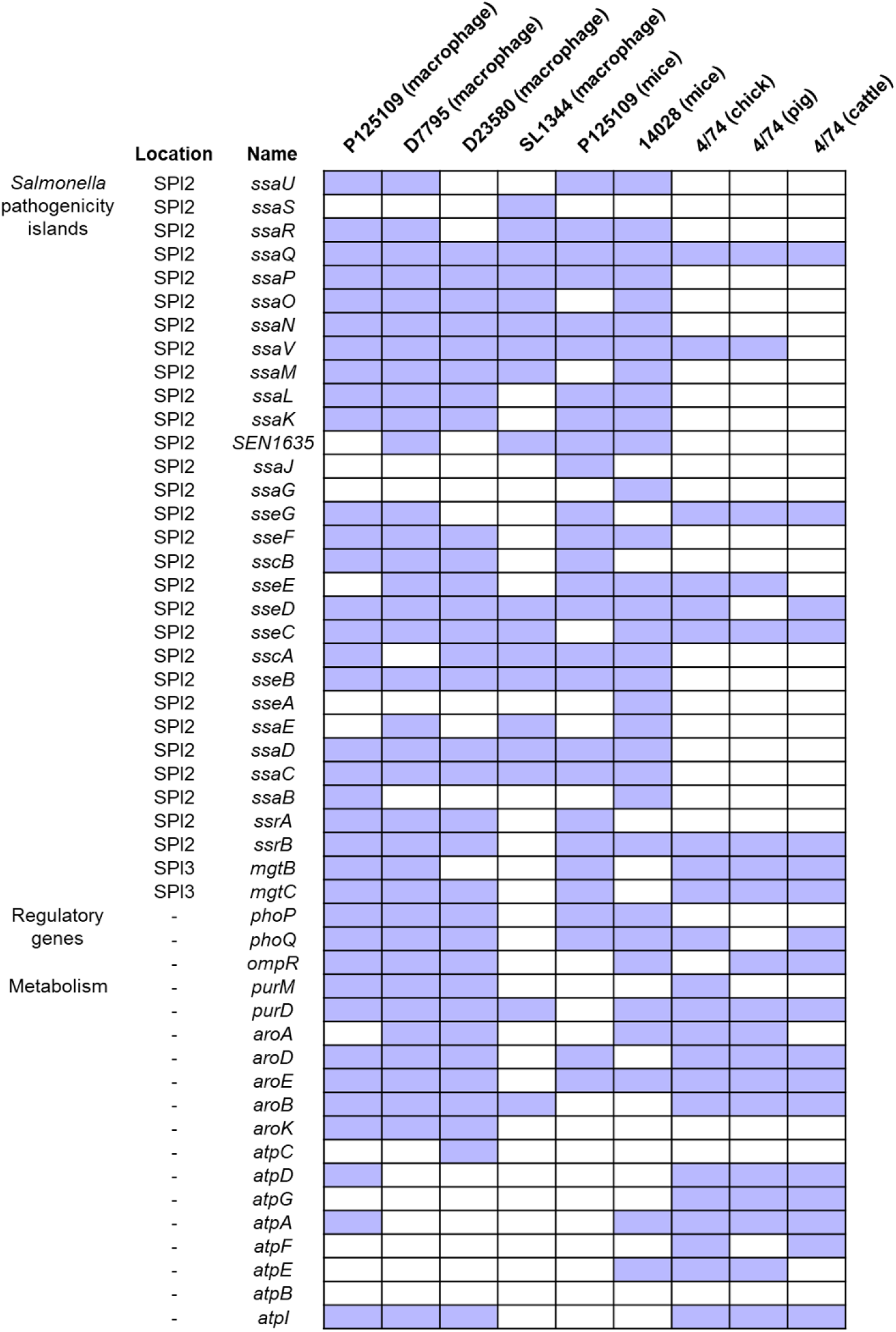
Macrophage-fitness genes in *S*. Enteritidis P125109 and D7795 with reported roles in other *Salmonella* infection models. The figure shows 49 genes that are required for macrophage fitness in *S*. Enteritidis P125109 and/or D7795 as identified in this study, and their reported roles in other *Salmonella* infection models. Blue box indicates the gene is involved in *Salmonella* fitness in the specified strain and infection model. D23580 (macrophage) = *S*. Typhimurium ST313 D23580 in macrophage infection [31]; SL1344 (macrophage) = *S*. Typhimurium ST19 SL1344 in macrophage infection [70]; P125109 (mice) = *S*. Enteritidis P125109 in BALB/c mice infection [16]; 14028 (mice) = *S*. Typhimurium ST19 14028 in BALB/c mice infection [73]; and 4/74 (chick), 4/74 (pig), 4/74 (cattle) = *S*. Typhimurium ST19 4/74 in food-related animal infection models [74].

**Table S1**. Bacterial strains and oligonucleotides used in this study.

**Table S2**. Illumina DNA libraries generated in this study and number of sequenced reads for each sample at every step.

**Table S3**. Number of reads, transposon insertion sites, insertion index, and essentiality call per gene. Samples included: P125109 input (for LB, NonSPI2 and InSPI2), P125109 LB (output), P125109 NonSPI2 (output), P125109 InSPI2 (output), P125109 input (macrophage), P125109 macrophage (output), P125109 LB_10h (output), D7795 input (for LB, NonSPI2 and InSPI2), D7795 LB (output), D7795 NonSPI2 (output), D7795 InSPI2 (output), D7795 input (macrophage), D7795 macrophage (output) and D7795 LB_10h (output).

**Table S4**. Raw data for figures.

**Table S5**. Analysis of the TIS macrophage data. Read counts for the two inputs and two outputs (LB_10h and macrophage), and log_2_ fold-changes and adjusted *P*-values for the comparative analysis of each coding gene and non-coding sRNA in *S*. Enteritidis P125109 and D7795.

**Table S6**. Comparison of macrophage-attenuated genes in *S*. Enteritidis GEC P125109 and CEAC D7795 with genes associated with virulence in other *Salmonella* serovars and infection models: *S*. Typhimurium ST313 D23580 [31] and ST19 SL1344 [70] in macrophage infection; P125109 [16] and *S*. Typhimurium ST19 14028 in BALB/c mice infection [73]; and *S*. Typhimurium ST19 4/74 in food-related animal infection models [74].

**Table S7**. Cluster of Orthologous Genes (COGs) categories assigned by eggNOG mapper

## Notes

https://tinyurl.com/GECP125109

https://tinyurl.com/CEACD7795

https://www.ebi.ac.uk/ena/browser/view/PRJEB52017

### Summary of Updates

Impact statement and new Figure S2 added; author current affiliations updated

