## Supplemental Table 7 for "Genome-wide fitness analysis identifies genes required for *in vitro* growth and macrophage infection by African and Global Epidemic pathovariants of *Salmonella* Enteritidis"

Table S7 Cluster of Orthologous Genes (COGs) categories assigned by eggNOG mapper

|  | **COG** | **Description** | **Example functions/proteins and associated genes** |
| --- | --- | --- | --- |
| **Information Storage and Processing** | J | Translation, ribosomal structure and biogenesis | tRNA-synthetases: *glnS*, *proS*, *ileS*, *leuS*  Ribosome components: *rpl* genes, *rpm* genes, *rps* genes  Initiation, elongation, and peptide chain release factors: *infA*, *efp*, *prfAH*, *pth*  RNA chaperones: *proQ*, *hfq* |
|  | A | RNA processing and modification | RNA 3’-terminal phosphate cyclase: *rtcA*  Oligoribonuclease (3’-5’ exoribonuclease): *orn* |
|  | K | Transcription | RNA polymerase: *rpoABC*  Sigma factors: *rpoESD*  Anti- and termination factors: *nusBAG*  Initiation, elongation, termination factors  Transcriptional regulators: *hns*, *fnr*, *hilA*  Nucleoid-associated proteins: *hns*  Response regulators of two-component systems (TCS): *phoP*  Cold shock proteins *csp* |
|  | L | Replication, recombination and repair | Primosome-associated: *dnaGAC*, *priABC*, *rep*, *ssb*  DNA polymerase III: *dnaEN*, *holABCE*  DNA polymerase II: *polB*  DNA polymerase I: *polA*  Gyrase and topoisomerases: *gyrAB*, *parCE*  Endo- and exonucleases: *mutHLMT*, *rnh*, *ruvAC*, *uvrABCD*  Transposase, integrase |
| **Cellular Processes and Signalling** | D | Cell cycle control, cell division, chromosome partitioning | Cell division proteins: *damX*, *ftsAEUKLNQWXYZ*, *sulA*  Chromosome partitioning: *mukBEF*  Cell partitioning: *minCD* |
|  | V | Defense mechanisms | Transporters and efflux pumps: *sapBC*, *acrDEF*  Restriction enzymes: *hsdMRS*  Beta-lactam-associated: *ampEDH* |
|  | T | Signal transduction mechanisms | Sensor kinases of TCS: *envZ*, *phoQ*, *arcB*, *baeS*, *ssrA* |
|  | M | Cell wall / membrane / envelope biogenesis | Peptidoglycan synthesis: *murABCDEFGI*, *mraY*  Lipopolysaccharide biosynthesis: *rfaBCFGIJKLQ*  O-antigen biosynthesis: *rfb* genes  Colanic acid polysaccharide capsule biosynthetic genes: *wca* genes |
|  | N | Cell motility | Fimbrial proteins: *bcfDEFG*, *fim*ADFHI, *pegABD, stbACD*  Flagella biosynthesis & components: *flgABCDEFGHIJKLMN*, *flhABE*, *fliCDEFGH*, *motAB*  SPI secretion system apparatus proteins: *spaOS* (SPI-1), *ssaCKNQ* (SPI-2) |
|  | W | Extracellular structures | Autotransporter adhesin: *sadA*  Outer membrane usher protein *fimD* |
|  | U | Intracellular trafficking, secretion, and vesicular transport | Sec translocase  SPI-1 TTSS (*spaPQR*)  SPI-2 TTSS (*ssaJRSTUV*) |
|  | O | Post-translational modification, protein turnover, chaperones | *ccm* genes  *clpABPX*  Molecular chaperones: *dnaAK*  Fe-S cluster: *sufBCD*  Lon protease: *lon*  *ppiBAC* |
| **Metabolism** | C | Energy production and conversion | ATP synthase: *atpBEFGH*  Cytochrome: *cybBC*, *cydABCD*  *nuo* genes |
|  | G | Carbohydrate transport and metabolism | Galactose metabolism and transport: *galKMPT*  Galactarate metabolism and transport: *garDKL*  Mannose metabolism and transport: *manAXYZ* |
|  | E | Amino acid transport and metabolism | Arginine biosynthesis: *argACDEH*,  Aromatic amino acids biosynthesis: *aroABCDEF* |
|  | F | Nucleotide transport and metabolism | Purine metabolism: *purABCDEFGH* |
|  | H | Coenzyme transport and metabolism | Heme biosynthesis: *hem* genes  Aspartate pathway: *nadABCDE*  *ribABDEF* genes |
|  | I | Lipid transport and metabolism | *accABCD*  Fatty acid biosynthesis: *fabABDFGH* |
|  | P | Inorganic ion transport and metabolism | *fhu* genes  Catalase genes*: kat* genes  Magnesium transport: *mgtABC* |
|  | Q | Secondary metabolites biosynthesis, transport and catabolism | Siderophore (enterobactin) synthesis: *entBCDEF* genes  *pduACEJKLNTX* (although some are also classified under category C) |
| **Poorly Characterised** | S | Function unknown | Includes many genes associated with Regions of Difference (RODs), prophage regions and SPIs |

Note: eggNOG-mapper uses only 20 of the 26 functional categories from the NCBI COG database (<https://www.ncbi.nlm.nih.gov/research/cog>). An “N/A” category was added to group all genes that were not assigned a COG category by eggNOG-mapper.
